# Predator coexistence and herbivore suppression are shaped by predator functional types in intraguild predation modules

**DOI:** 10.64898/2026.05.21.726930

**Authors:** Emilio Mora Van Cauwelaert, Enric Frago, Francisco Martínez-Martínez, Vasilis Dakos

**Affiliations:** Institut des Sciences de l’Évolution de Montpellier, Universite de Montpellier (ISEM), CNRS, Montpellier, France; CIRAD, UMR CBGP, INRAE, Institut Agro, IRD, Université Montpellier, Montpellier, France

**Keywords:** intraguild predation, coexistence, functional types, biocontrol

## Abstract

Coexistence of multiple predators in ecological communities and their combined effects on the abundance and diversity of shared prey are often difficult to predict. In some theoretical models, predator coexistence is limited by antagonistic interactions, especially in the form of intraguild predation (IGP) that typically leads to out-competition between predators and high prey densities. However, empirical studies show that predator coexistence is common even in the presence of IGP. This discrepancy between theoretical expectations and empirical observations can be highly relevant for practical applications like using multiple natural enemies for pest suppression in agriculture. It is proposed that greater functional differences between natural enemies (i.e. predators) could reduce competition and overcome the negative effects of IGP, thereby promoting their coexistence and enhancing herbivore (i.e. prey) suppression. In this study, we theoretically explore this proposition. We develop a theoretical model based on the types of natural enemies of aphids to identify how functional differences between predators in IGP modules affect predator coexistence and herbivore suppression. We show that pairwise combinations of four functional predator types (ladybird, predatory bug, hoverfly, and parasitoid) can increase the coexistence range for different intraguild predation and competition strengths between predators (IGP symmetry), along a productivity gradient. This outcome depends on the external food input rate for the predatory bug and hoverfly types, and on their position as IG predator or IG prey. Herbivore suppression was primarily driven by IGP symmetry (i.e. the relative intraguild predation and exploitative competition strength between predators) and was especially pronounced in competitive-like modules where the IG predator was excluded for most scenarios. However, for some competitive-like IGP modules with predatory bug and hoverfly types, both predators can persist and provide a high herbivore suppression across increasing productivity. Our results can help explain experimental findings in conservation biocontrol, where coexistence between natural enemies is joined with effective herbivore suppression, and offer additional support for the role of functional diversity in reconciling theoretical predictions with experimental observations in multiple-predator communities.

## 1 INTRODUCTION

In ecological communities, predators are embedded in complex networks of interactions that include competition, facilitation, behavioral avoidance, and predation (McCoy et al., 2012). These interactions shape key ecological processes such as the likelihood of predator species to coexist in a community, or their impact on the abundance and diversity of species in lower trophic levels (Schmitz, 2007; McCoy et al., 2012). The study of multi-predator networks has yielded important theoretical insights regarding the role of higher-order interactions in the dynamics and stability of food webs (Mougi, 2022). It has also informed practical applications in agriculture, such as the use of multiple natural enemies to regulate the density of pest species (usually herbivores) (Oerke, 2006; Hajek and Eilenberg, 2018). In fact, one of the challenges of conservation biocontrol is the long-term persistence of natural enemies, while maintaining potential pests at low densities (Snyder, 2019; Landis et al., 2000; Hajek and Eilenberg, 2018). To cope with the complexity of analyzing multi-predator dynamics, some authors have focused on ecological “modules” rather than on complete communities (Polis et al., 1989). An ecological module is a community of a few species (usually 2 - 4) whose mechanisms and dynamics are relatively easy to model, yet sufficiently complex and realistic to represent the main structures and patterns observed in nature (Polis et al., 1989). One classical example is the Intraguild Predation module (IGP, Fig. 1A), where two predators that share the same prey can also feed on each other (the intraguild predation *per se*) (Polis et al., 1989; Arim and Marquet, 2004). Species in IGP modules are directly affected by predation, but they also experience competition and indirect mutualism interactions, resulting in very rich dynamics (Rosenheim and Harmon, 2006; Polis et al., 1989; Diehl and Feißel, 2000) (Figure 1A). These modules appear to be widespread in animal communities, and have been particularly studied in insect pests (Rosenheim et al., 1995).

**Figure 1:**
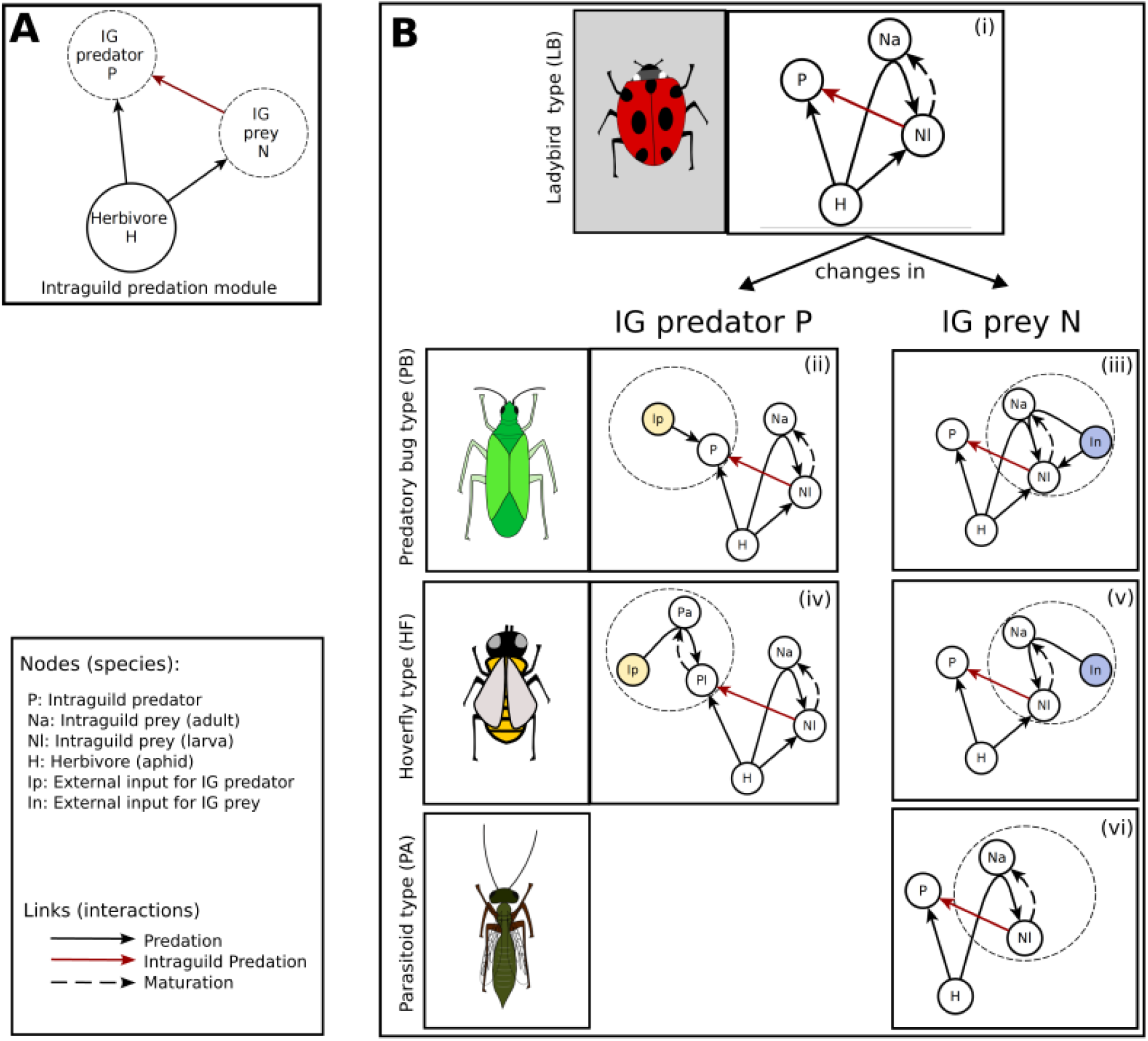
Construction of different Intraguild Predation Modules (IGP modules) combining four different predator typesA. The basic structure of the IGP module. Nodes represent species and arrows interactions between species. **B**. On the top, our baseline **LBLB** module where both IG predator *P* and IG prey *N* belong to the ladybird (LB) type (i). Herbivore *H* can be eaten by the IG prey larva (*N*_*l*_) and adult (*N*_*a*_), as well as by the IG predator *P*. Predation from *N*_*a*_ directly transforms to biomass of the larva *N*_*l*_. The IG predator also preys on the IG prey larva but not on the IG prey adult. *N*_*l*_ matures into *N*_*a*_. Lower insets show the changes incurred to the top baseline module when other predator types are considered, either as IG predator *P* or as IG prey *N* (Table 1). The big dotted circles highlight the differences to the baseline **LBLB** module. When the Predatory Bug type (PB) is considered as IG predator *P* or as IG prey *N*, we add external inputs *I*_*P*_ (yellow) to *P* or *I*_*N*_ (blue) to both *N*_*a*_ and *N*_*l*_ (ii and iii). When the Hoverfly type (HF) is considered as IG predator *P*, the IG predator *P* is structured into adults *P*_*a*_ and larvae *P*_*l*_ and the external food input *I*_*P*_ is consumed only by the adult *P*_*a*_ (iv). By symmetry, when the HF type is considered as an IG prey, the external food input *I*_*N*_ is consumed only by adults *N*_*a*_ (v). The parasitoid type (PA) is considered only as an IG prey (and not IG predator). The infected larvae *N*_*l*_ of the PA type lie within the herbivore and is promoted by the infection of the *N*_*a*_ of the herbivore *H* (vi). The drawings are made by E. Frago.

According to theoretical models of IGP, coexistence among intraguild predators is unlikely, it depends on the availability of resources (or the productivity of the system), and it always leads to a higher density of prey than without coexistence (Rosenheim and Harmon, 2006; Polis et al., 1989; Diehl and Feißel, 2000; Sentis et al., 2013). However, experimental and field studies show that coexistence is ubiquitous in multi-predator communities, and it does not always lead to high prey densities but also to prey suppression (Chang and Cardinale, 2020; Rosenheim and Harmon, 2006). This discrepancy between theoretical expectations and empirical observations could be explained by the fact that theoretical predictions mostly come from simple models that do not account for more realistic conditions like spatial dynamics, seasonality or behavioral changes(Brodeur and Boivin, 2006). Early additions to IGP models included some of these complexities such as stage structure (Mylius et al., 2001), temporal refugees (Amarasekare, 2008), alternative prey (Holt and Huxel, 2007; Briggs and Borer, 2005), or adaptive predation and behavior (Krivan and Schmitz, 2003; Hentley et al., 2016). Yet, although most of these expanded models increase the likelihood of predator coexistence, they still uphold the basic prediction that predator coexistence reduces the suppression of the shared prey (Rosenheim and Harmon, 2006) (but see (Briggs and Borer, 2005)). Specifically, most models show that, for coexistence to be achieved, the eaten predator (the intraguild prey) must be a better competitor for the shared prey, and its reduction in the presence of the other predator (the intraguild predator) “releases” the pressure on the prey, increasing its density (Polis et al., 1989; Diehl and Feißel, 2000). The population of the shared prey is also affected by the relative feeding of the intraguild predator on the intraguild prey and on the shared prey (Faria and Costa, 2009; Vandermeer, 2006; Krivan and Diehl, 2005). This prey-preference of the intraguild predator (*the IGP symmetry*) can modify the impact of intraguild predation on the competition between the two predators for the shared prey and on the eventual control of the prey population.

**Table 1:** Most common predator types involved in an Intraguild Predation (IGP) module. The classification is based on their predatory stage-structure, the presence or absence of an alternative diet, and their mode of intraguild predation (coincidental or intentional). We named the predator types according to a representative predator of aphids that belonged to that predator type but are not exclusive to that predator species. For example, the Ladybird type (LB) includes ladybirds (Coleoptera) and species of brown lacewings (Neuroptera). (*) Both in LB and PA types, some adults complement their diet with pollen or nectar, but it is not mandatory for survival.

| Predator type | Predatory stage structure | Alternative diet | Mode of intraguild predation | Examples |
| --- | --- | --- | --- | --- |
| **Ladybird type (LB)** | Larvae and adults are predatory | No mandatory external source of food (*) | Intentional | Predatory ladybirds (Coleoptera), brown lacewings (Neuroptera) |
| **Predatory Bug type (PB)** | Larvae and adults are predatory | Both stages feed on external insects, pollen and nectar | Intentional | Predatory bugs (Hemiptera) |
| **Hoverfly type (HF)** | Only larvae are predatory | Adults feed exclusively on nectar and pollen | Intentional | Hoverflies and predatory flies (Diptera) |
| **Parasitoid type (PA)** | Adults oviposit on hosts and larvae grow within the host | No mandatory external source of food (*) | Coincidental | Braconid, figitid and encyrtid wasps (Hymenoptera) |

However, some studies have proposed that complementarity or resource partitioning between predators that share the same prey could reduce the impact of interspecific competition and promote their coexistence (Snyder, 2019). Complementarity in predator assemblages is often determined by the degree of functional dissimilarity between species, which includes their life cycle, their degree of diet specialization, or their hunting strategy (Alhadidi et al., 2019; Schmitz, 2007). For example, a predator that undergoes substantial diet shifts during its life cycle or has alternative sources of food is likely to lower overall competition to a competing predator with another diet regime (Mylius et al., 2001; Hin et al., 2011; Rudolf and Lafferty, 2011). This would imply that functional dissimilarity could allow for coexistence between predators without necessarily increasing the population density of the shared prey (Snyder, 2019; Alhadidi et al., 2019; Chang and Cardinale, 2020). Yet, it is still not fully understood how functional differences modulate co-existence and prey suppression in IGP modules. In addition, it is not clear how these differences interact with IGP modules with different IGP symmetries (from more tritrophic to competitive-like modules) and along a gradient of productivity (but see (Diehl and Feißel, 2000; Sentis et al., 2013).

Understanding the conditions for predators’ coexistence and their joined effect on shared prey is particularly relevant in the context of biocontrol in agriculture (Alhadidi et al., 2019; Brodeur and Boivin, 2006; Perović et al., 2018). Predators used for herbivore suppression (termed natural enemies) are exemplary cases of intraguild predation behavior (Rosenheim et al., 1995). Natural enemies are usually very diverse and many functional groups are commonly associated to any given pest, their diversity depending on the ecological context, the herbivore targeted, and the cultivated plants (Snyder and Tylianakis, 2012). In particular, IGP interactions have been intensively studied in aphids (Aphididae), not only because they are an important group of pests, but also because they have a very diverse list of natural enemies, including more than six biological orders and many distinct functional groups (Alhadidi et al., 2019; Vance-Chalcraft et al., 2007; Sentis et al., 2013; Hentley et al., 2016). These can be specialists of aphids (e.g. ladybirds) or feed also on external food sources such as nectar, pollen, plants or other insects (e.g. syrphids or predatory bugs) (Daugherty et al., 2007; Van Rijn et al., 2002). They can differ on whether they change their diet during their life cycle (e.g. syrphids) or not (e.g. ladybirds) (Mylius et al., 2001). Natural enemies can also be intentionally or coincidentally eaten by other predators (Rosenheim et al., 1995). In some cases, predators’ immature stages are directly preyed upon, but in others, their larvae lie within an infected herbivore and are coincidentally eaten (e.g. parasitoids). Therefore, as IGP modules of natural enemies of aphids can be composed by different combinations of these functionally different predators, they constitute a perfect model system to theoretically explore the influence of functional dissimilarity on coexistence and prey suppression. This system is also a good model as the functional types of predators can be extrapolated to most arthropods groups. Experimental studies have tested some combinations of functionally different natural enemies of aphids (predators vs parasitoids, generalist vs specialist), showing non-consistent effects on short term prey suppression which suggest that more realistic and mechanistic models are needed (Alhadidi et al., 2019).

In this paper, we proposed a mathematical model mimicking relevant empirical setups of natural enemies to identify how IGP modules composed of predators from different functional groups affect their probability of coexistence and modify herbivore suppression. We defined four functional predator “types” based on the natural enemies of aphids, and modeled how combinations of these types modified their coexistence and the level of herbivore suppression in IGP modules. We also explored how these combinations interacted with different levels of IGP symmetry (relative intraguild predation and exploitative competition strength) along a gradient of resource productivity. We discussed then the mechanisms behind coexistence and herbivore suppression according to functional dissimilarity of predators, compare our theoretical expectations to empirical observations, and outline implications of our findings in agricultural systems (Perović et al., 2018; Diehl et al., 2013).

## 2 MODEL AND METHODS

### 2.1 Predator types and IGP modules

We classified natural enemies of herbivores into four “predator types” according to the following life history traits: (a) diet changes through their life cycle, notably between larval or nymph stages and adults (“predatory stage structure”), (b) necessity to feed or not on external sources of food such as pollen or plants (“alternative diet”), and (c) the way natural enemies suffer predation within an IGP module, either because their nymph or immature stage is intentionally eaten by the other predator, or if it occurs coincidentally (sensu (Rosenheim and Harmon, 2006)) because the larvae lie within a potentially eaten infected herbivore (“mode of intraguild predation”). Based on these three traits using aphid predators as case study, we defined four predator types (Hajek and Eilenberg, 2018): the LadyBird type (LB), the Predatory Bug type (PB), the Hover- Fly type (HF), and the Parasitoid type (PA) (Table 1). In the case of aphids, the **Ladybird type (LB)** contains specialist predators of aphids where both the larvae and the adults are predatory (i.e., no changes in diet across stage structure) and that do not need external sources of food (for most species, feeding on external sources is possible as a complement, but not mandatory for survival). This type includes predatory ladybirds (Coleoptera: Coccinellidae), but also most species of brown lacewings (Neuroptera: Hemerobiidae). The **Predatory Bug type (PB)** includes generalist predators that also feed on external sources of food, usually of plant origin, and where both the adults and the larvae are predatory (no changes in diet across stage structure). This category includes many species from different groups, mostly predatory bugs in the order Hemiptera. The **Hoverfly type (HF)** includes predators where only the larvae feed on aphids, and the adults have other sources of food, usually nectar or pollen. This predator type includes hoverflies (Diptera: Syrphidae) and some predatory flies like aphid midges (Diptera: Cecidomyiidae). In the **Parasitoid type (PA)**, adults lay their eggs inside aphids, and the internally hatched larvae feed on the aphid and can be coincidentally eaten by another predator. Aphid parasitoids are highly specialized in aphids, particularly certain groups of braconid wasps (Hymenoptera: Braconidae), but also less specialized wasps like figitids (Hymenoptera: Figitidae) and encyrtids (Hymenoptera: Encyrtidae). Our classification is based on the natural enemies of aphids, but it can be easily extended to other pests and their associated predators, because the three predatory traits we used can be considered in any other predator-prey system (Hajek and Eilenberg, 2018). For example, predatory bug and parasitoid types are universal groups of natural enemies present for most prey groups; praying mantis, carabid beetles or predatory mites could be considered as ladybird types; and any predator group in which adults are pollen feeders -like fireflies, net-winged beetles, antlions, and owlflies- can be considered as hoverfly type predators (Hajek and Eilenberg, 2018).

### 2.2 IGP model of the baseline ladybird-ladybird module (LBLB)

In our system, IGP modules of natural enemies are made up of pairs of predators from different predator types. An IGP module includes an intraguild predator (IG predator) that feeds both on the herbivore and on the intraguild prey (IG prey), which also eats the herbivore (Fig. 1A). The ladybird-ladybird module (**LBLB**)) was formed by predators that belonged to the ladybird (LB) predatory type (Table 1). The model was based on the IGP model of Polis et al. (1989) with stage structure of the IG prey as in Mylius et al. (2001) (Fig. 1B). This stage-structure allowed us to represent how different natural enemies can feed on one another as they attack the larval stage which tends to be smaller. We consider the **LBLB** module as the baseline module because both natural enemies belonged to the LB type, which represents the functionally simplest predator as it does not feed on any external sources of food and does not have a different diet across its developmental stages. The model of the **LBLB** module included a herbivore *H* (i.e. the pest), an IG prey *N* with larval and adult stages *N*_*l*_ and *N*_*a*_ respectively, and an IG predator *P*.

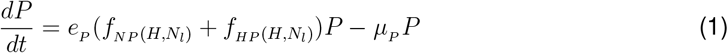

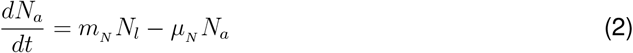

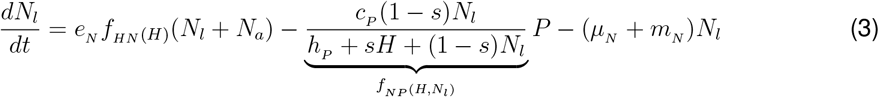

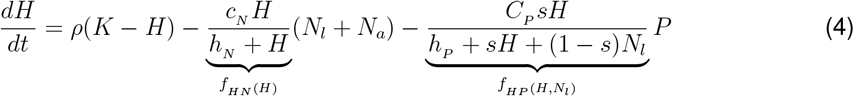

The herbivore *H* (eq.4) follows semi-chemostat dynamics in the absence of predators, with turnover *ρ* and carrying capacity *K* (representing the productivity of the system). *H* can be eaten by the IG prey larvae *N*_*l*_ (eq. 3) and IG prey adult *N*_*a*_ (eq. 2), as well as by the IG predator *P* (eq. 1) with functional responses 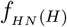 and 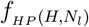. This is converted into biomass with energy efficiencies *e*_*N*_ and *e*_*P*_. Both predators also have a density-dependent death rate *µ*_*N*_ and *µ*_*P*_. The IG predator *P* also preys on *N*_*l*_ with functional response 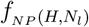. The larva *N*_*l*_ transforms into an invulnerable adult *N*_*a*_ at a *m*_*N*_ rate. Importantly, we assumed that all consumption of adults *N*_*a*_ directly transforms to biomass of the larvae *N*_*l*_. We used saturating type II functional responses with half saturation constants *h*_*N*_ and *h*_*P*_ and maximum ingestion rates *c*_*N*_ and *c*_*P*_. This type of functional response accounts for the provoked scarcity of insects and is a good predictor for IGP dynamics with changing productivities (Sentis et al., 2013). Finally, we included a parameter *s* that changes the relative feeding of the IG predator *P* from a tritrophic-like food chain (when *s ≈* 0, *P* feeds mainly on *N*), to a symmetric IGP (when *s ≈* 0.5, *P* feeds equally on *N* and on *H*), to an exploitative competition-like module (when *s ≈* 1, *P* only feeds on *H*). This parameter of symmetry regulates the relative strength of the intraguild predation and the exploitative competition between the predators on the shared herbivore resource (Faria and Costa, 2009; Vandermeer, 2006).

### 2.3 Modeling different predator types in IGP modules

In order to build the rest of the IGP modules with different predator types, we modified the interactions and the equations of the baseline **LBLB** model (Fig. 1B and Fig. S.1 for all IGP modules structures). Contrary to the Ladybird type (LB), the Predatory bug type (PB) includes generalist predators that feed on external food sources (for example, other insects or plants, Table 1). This was modeled as an external food input *I*_*P*_ for the IG predator *P*, or as input *I*_*N*_ for both adult *N*_*a*_ and larva *N*_*l*_ for the IG prey *N* (yellow and blue nodes in Fig. 1B, respectively). The Hoverfly type (HF) includes predators where only the larvae feed on the herbivore whereas the adults have other sources of food, usually nectar or pollen (Table 1). We modeled this by dividing the IG predator *P* into an adult *P*_*a*_ and a larval *P*_*l*_ stage, where only *P*_*l*_ was predatory and where *P*_*a*_ fed only on an external input *I*_*P*_. We did the same also when HF type was an IG prey *N* with an external input *I*_*N*_ (yellow and blue nodes Fig. 1B). In the case of the Parasitoid type (PA), adults parasitized the herbivore and the internally hatched larvae fed on the infected herbivore (Table 1). We modelled the parasitoid as an IG prey *N* assuming that the adult *N*_*a*_ infects the herbivore *H* to produce a larva *N*_*l*_ that does not eat the herbivore, but grows into it. We also assumed that *N*_*l*_ does not have an internal mortality rate as it lies within the herbivore, and transforms to an adult *N*_*a*_ (Fig. 1, see Nakazawa et al. (2010)). We did not add the parasitoid type as an IG predator *P*, as we did not consider cases of hyper-parasitism (but see Schreiber et al. (2001)). For all models, external food *I*_*P*_ and *I*_*N*_ was added as constant inputs rates whose consumption was density dependent (Brodeur and Boivin, 2006). For predators belonging to the Predatory bug type (PB) we added preferences *ϕ*_*P*_ and *ϕ*_*N*_ that modulate the relative feeding of the external input. In total, we combined three predator types (LB, PB, HF) as IG predators *P* with four predator types as IG prey *N* (LB, PB, HF, PA) resulting in 12 IGP combinations, all represented by a set of ordinary differential equations detailed in section 1 of the Supplementary Material.

### 2.4 Model parameterization

We parameterized the baseline **LBLB** module following Hin et al. (2011) (Table 2). In their work, the IG prey’s maintenance rate is fixed to 1, and the other parameters are scaled through body-mass relationships (Hin et al., 2011). Here, we included the maintenance rate of each predator within their death rate. We confirmed that parameter values satisfied the Polis and Holt (1989) *necessary* criterium for coexistence for any value of IGP symmetry *s* and productivity *K*. This criterium states that the IG prey *N* needs to be a superior competitor than the IG predator *P* for the shared herbivore (section 2 in the Supplementary Material). To allow comparisons between the different IGP modules, we used the same set of parameters. For the Predatory bug (PB) or the Hoverfly (HF) types, as IG predators or IG prey, we calculated the maximum rate of the external inputs (*I*_*P*_ or *I*_*N*_) above which the predator could live only on the external input (given our assumption that larvae have a density independent maturation rate). These values depended on the predator type and on whether the predator was the IG prey *N* or IG predator *P* (section 2 in the Supplementary Material). Next, to ensure that our module could be defined as an IGP system (i.e. both predators are dependent on the herbivore), we took the 10 and 90% of this maximum rate and labeled them as low and high external input (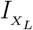 and 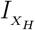 where *X ∈ {N, P}*, Table 2). Finally, for the scenarios with the generalist Predatory bug (PB) type we assumed that both larval and adult stages have an equal and non adaptive preference for the external input (*ϕ* = 0.5 of a scale from 0 to 1). This decision also helped us to standardize the total consumption of predators that fed on external food to those that did not, across IGP modules. We also ran sensibility tests for other values of *ϕ*_*P*_ and *ϕ*_*N*_ which did not qualitatively changed the results (section 3 in the Supplementary Material).

**Table 2:** Parameters, values and units and their definitions used for all 12 IGP modules. For all variables *P*, *N*_*l*_, *H, I*_*P*_ and *I*_*N*_, biomass units are qualitatively equivalent. The time is written in time units (*t*.*u*,). Note that when HF is the predator type, we have to change *P* to *P*_*l*_ in *C*_*P*_ and *e*_*P*_ units, and *P* to *P*_*a*_ in *I*_*P*_ units.

| Parameter | Value | Unit | Definition |
| --- | --- | --- | --- |
| **IG predator $P$** |  |  |  |
| $C_P$ | 2.5 | $(t.u.)^{-1}$ | Maximum ingestion rate |
| $h_P$ | 1 | biomass$_{(H+N_L)}$ | Half saturation constant |
| $e_P$ | 0.5 | — | Conversion efficiency |
| $\mu_P$ | 0.33 | $(t.u.)^{-1}$ | Natural death rate |
| $m_P(HF)$ | 0.2 | $(t.u.)^{-1}$ | Maturation rate when HF |
| $I_P(PB)$ | {0.13, 1.2} | $(t.u.)^{-1}$ | External input rate when PB |
| $I_P(HF)$ | {0.17, 1.6} | $(t.u.)^{-1}$ | External input rate when HF |
| $\phi_P(PB)$ | 0.5 | — | External input preference when PB |
| $s$ | [0 – 1] | — | IGP symmetry (relative feeding on H or N) |
| **IG prey $N$** |  |  |  |
| $C_N$ | 10 | $(t.u.)^{-1}$ | Maximum ingestion rate |
| $h_N$ | 1 | biomass$_{(H)}$ | Half saturation constant |
| $e_N$ | 0.5 | — | Conversion efficiency |
| $\mu_N$ | 1.1 | $(t.u.)^{-1}$ | Natural death rate |
| $m_N$ | 0.2 | $(t.u.)^{-1}$ | Maturation rate |
| $I_N(PB)$ | {0.44, 4} | $(t.u.)^{-1}$ | External input rate when PB |
| $I_N(HF)$ | {1.4, 13} | $(t.u.)^{-1}$ | External input rate when HF |
| $\phi_N(PB)$ | 0.5 | — | External input preference when PB |
| **Herbivore $H$** |  |  |  |
| $\rho$ | 1 | $(t.u.)^{-1}$ | Chemostat turnover |
| $K$ | [0 – 8] | biomass$_{(H)}$ | Productivity of the system |

### 2.5 Assessing species coexistence and herbivore suppression

We assessed coexistence of both IG predator *P* and IG prey *N* for each of the 12 IGP modules by performing a 2-D bifurcation analysis for a gradient of IGP symmetry (*s* ranging from 0 to 1 in steps of 0.01), and of productivity (*K* ranging from 0 to 8 in steps of 0.01; the maximum productivity was chosen to include all the qualitative behaviors of all IGP modules). For each combination of productivity and IGP symmetry, we numerically solved for the asymptotic equilibria by simulating for 2000 time steps. For the IGP modules with the PB and HF types, we analyzed two scenarios for low and high values of the external input (Table 2). To evaluate the effect of the different predator types on coexistence and herbivore suppression, we used the **LBLB** module as baseline and compared the coexistence region (where both predators were present along the 2-D bifurcation plot) to the coexistence region of all other IGP modules. To evaluate the impact of the IGP modules on herbivore suppression we assessed whether herbivore density was reduced relative to the **LBLB** baseline module. For this, we extracted and compared for each IGP module, the equilibrium herbivore densities along the productivity gradient, for three values of IGP symmetries *s* that represented tritrophic-like (*s* = 0.1), symmetric (*s* = 0.5) and competitive-like (*s* = 0.9) modules. For the IGP modules where PB or HF acted as IG predator *P* or as IG prey *N*, we used the values of external inputs *I* and *I* where coexistence was enhanced (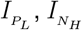). We performed these comparisons both when coexistence of predators was possible or when not. Bifurcation analysis was performed with the R *debif* package (version 0.1.9) (de Roos, 2025). We confirmed results of the bifurcation analysis by numerical simulations to identify the different equilibria and the transition boundaries between them. We did this by simulating the different models for 2000 time steps (by 0.05 integration step) starting from randomized initial conditions and identifying asymptotic equilibria with a tolerance threshold of 10^−6^ using the deSolve package with LSODA. All analyses were implemented in RStudio 2023.03.1. The documented code, the used libraries, and data required to reproduce results can be accessed at https://github.com/tenayuco/igp-predatorTypes.

## 3 RESULTS

### 3.1 Equilibria and Coexistence in the baseline LBLB IGP module

The **LBLB** module represented the baseline IGP module with the simplest structure where both predators belonged to the Ladybird type LB. We identified seven stability regions along the two-dimensional axes of IGP symmetry *s* and productivity *K*, with four equilibria possible as found elsewhere (Mylius et al., 2001): (1) a herbivore-only equilibrium (*H*^*^), (2) an IG prey *N* - herbivore *H* equilibrium (*HN*^*^), (3) a coexistence equilibrium where the three species are present (*HNP*^*^) and (4) an IG predator *P* -herbivore *H* equilibrium (*HP*^*^).

The coexistence region –defined as the region in the 2-D bifurcation plot of *s* and *K* where both the IG prey *N* and the IG predator *P* were present (*HNP*^*^)– was restricted to medium and low values of IGP symmetry (*s <* 0.6) and to sufficiently high productivities (*K >* 1) (green zone in Fig. 2A). In this region, the IGP module satisfied a “mutual invasibility criterion” which states that given an IGP symmetry *s*, there are values of productivity *K* for which both *P* and *N* can invade an equilibrium where the other predator is present alongside the shared herbivore *H* (Supplementary Material S1). Given a highly tritrophic-like IGP module (*s* → 0), the coexistence equilibrium was stable in a wide range of productivity, and at high productivity levels (*K >* 6) the system crossed a Hopf bifurcation where the coexistence equilibrium became periodic (dark green zone in Fig. 2A). For symmetric IGP (*s ≈* 0.5), coexistence was restricted to a small range of productivity and allowed for the emergence of bistability of *HP*^*^ and *HNP*^*^ equilibria (light green zone in Fig. 2A). Above the region of bistability (*K >* 1.6) the IG predator *P* could invade and exclude the IG prey *N* (dark gray region in Fig. 2A). When the IGP modules were very competitive-like (*s* → 1), the IG prey *N* excluded the IG predator *P* even at high productivities (light gray region in Fig. 2A). The system also presented a zone of bistability of *HN*^*^ and *HP*^*^ equilibria, where none of the predators could invade the system if the other predator was present and where the final state depended on the initial densities of the competing predators (middle-gray zone in Fig. 2A). Finally, for low values of *K*, only the herbivore *H* was present (white region in Fig. 2), or the IG prey *N* could invade (light gray region in Fig. 2A).

**Figure 2:**
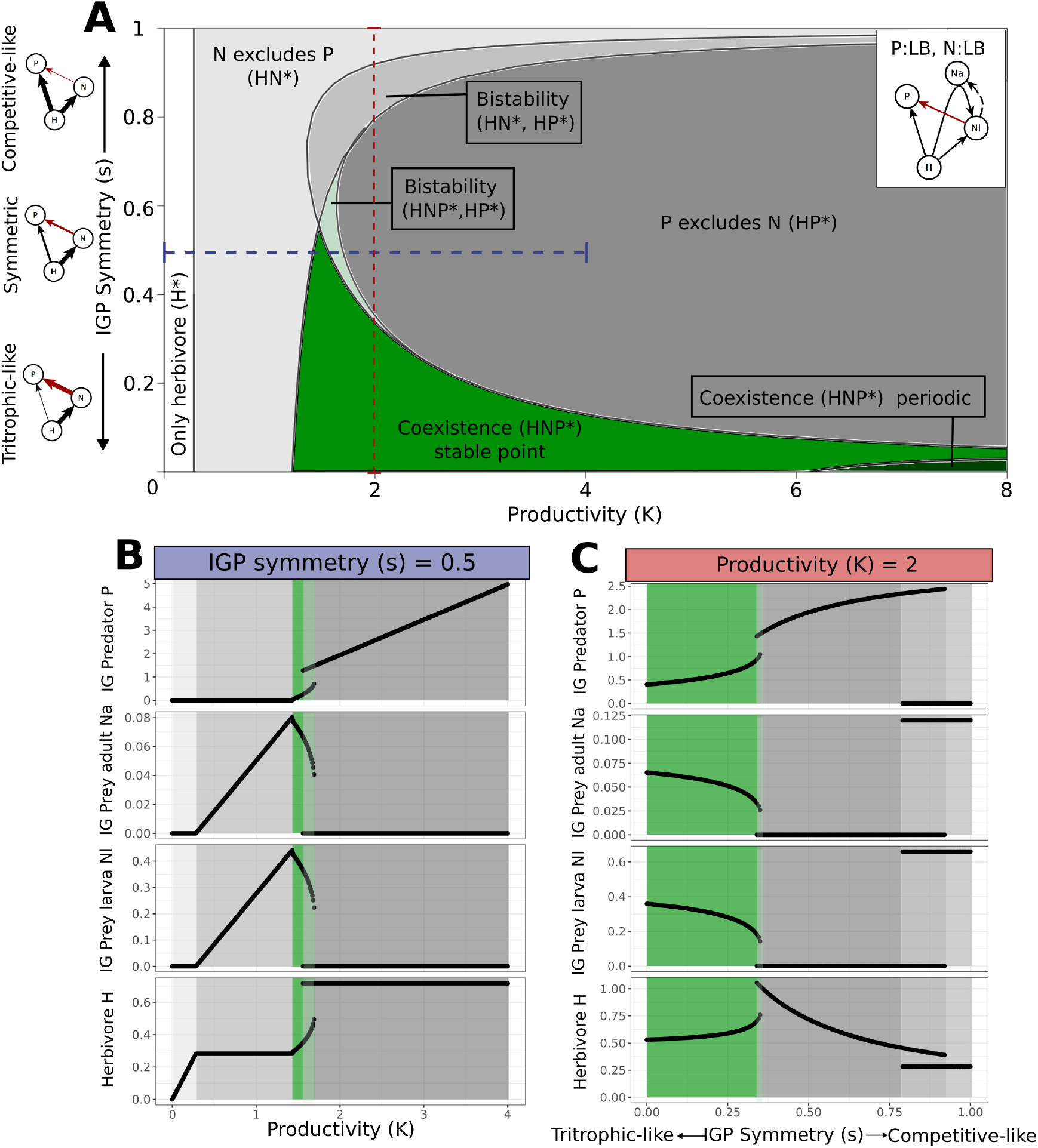
Equilibria and dynamics of the ladybird x ladybird IGP baseline module (LBLB). A. Asymptotic equilibria of the **LBLB** module across changing productivities *K* and IGP Symmetry values *s* ranging from no IGP (tritrophic-like, *s* = 0), to symmetric IGP (*s* = 0.5) and to no IGP again (*s* = 1, exploitative competition-like). Colors depict different equilibria: herbivore only equilibrium (*H*^*^, white), equilibria with one of the two predators only present (*HN* ^*^, light gray) or (*HP* ^*^, dark gray), bistability with one predator present(*HN* ^*^, *HP* ^*^, gray). Coexistence (*HNP* ^*^) was found either as stable fixed point equilibrium (green), periodic equilibria (dark green), or in a bistability region (*HNP* ^*^, *HP* ^*^, light green). For the equilibria representations we use *N* = *N*_*a*_ + *N*_*l*_. Straight dotted lines represent transects showing changes in population density of all species at equilibrium for constant *s* (panel B) and constant *K* (panel C). **B**. Population densities of the species at equilibrium along a gradient of productivity *K* and fixed *s* = 0.5 (blue) and **C** along a gradient of IGP symmetry *s* and a fixed *K* = 2 (red). Colors match the equilibria found in the 2-D bifurcation of panel A.

Given a symmetric IGP module (*s* = 0.5, Fig. 2B), the herbivore *H* persisted only with the IG prey *N* when productivity was low (*K <* 0.8, Fig. 2B). Increasing productivity *K* increased the IG prey equilibrium densities (both *N*_*l*_ and *N*_*a*_), while the herbivore density remained constant (balancing predation by the IG prey and its own carrying capacity). At medium productivity *K* (*K ≈* 1.5, Fig. 2B), a higher IG prey *N* density allowed the IG predator *P* to invade, causing a decrease in *N* and an increase in the herbivore (released from predation by *N*). For higher *K* (*K >* 1.8, Fig. 2B), a higher herbivore density supported the IG predator *P* who excluded the IG prey *N*, and the herbivore density remained constant again despite the increase in productivity. When productivity was fixed and intermediate (*K* = 2), increasing IGP-symmetry *s* made the system transition through all equilibria with at least one predator present (Fig. 2C). In the coexistence region, the IG prey (both *N*_*l*_ and *N*_*a*_) decreased with increasing *s*, while the IG predator *P* and herbivore *H* densities increased. Once the IG prey *N* was excluded (*s >* 0.3), a higher symmetry *s* increased the IG predator *P* pressure on the herbivore, reducing *H*. When the herbivore equilibrium density became too low, it did not support the IG predator *P* and allowed the IG prey *N* to invade and exclude *P*. In the absence of *P*, further increases in symmetry *s* did not affect the herbivore-IG prey *N* equilibrium. Between these last two regions lay a bistability region were both *HP*^*^ and *HN*^*^ equilibria were possible.

### 3.2 Coexistence across IGP modules with different predator types

Using as a baseline the **LBLB** module, we compared how coexistence changed across all other IGP modules composed by functionally different predator types (Table 1). In particular, we explored differences in the region of coexistence (within the 2-D bifurcation plot of *s* and *K*) by changing the IG predator *P*, the IG prey *N*, or both (Fig.3 and Fig. S1. for all the IGP structures). When the predator types could feed on the external inputs (*I*_*P*_ and *I*_*N*_), we included scenarios with low and high rates of the input. In addition, we described the population equilibria of the species across the new regions of coexistence of some representative IGP modules that maximized coexistence (Fig. 4), using a productivity value (*K* = 4) that encompassed the different qualitative behaviors of predators (dotted lines in Fig. 3).

**Figure 3:**
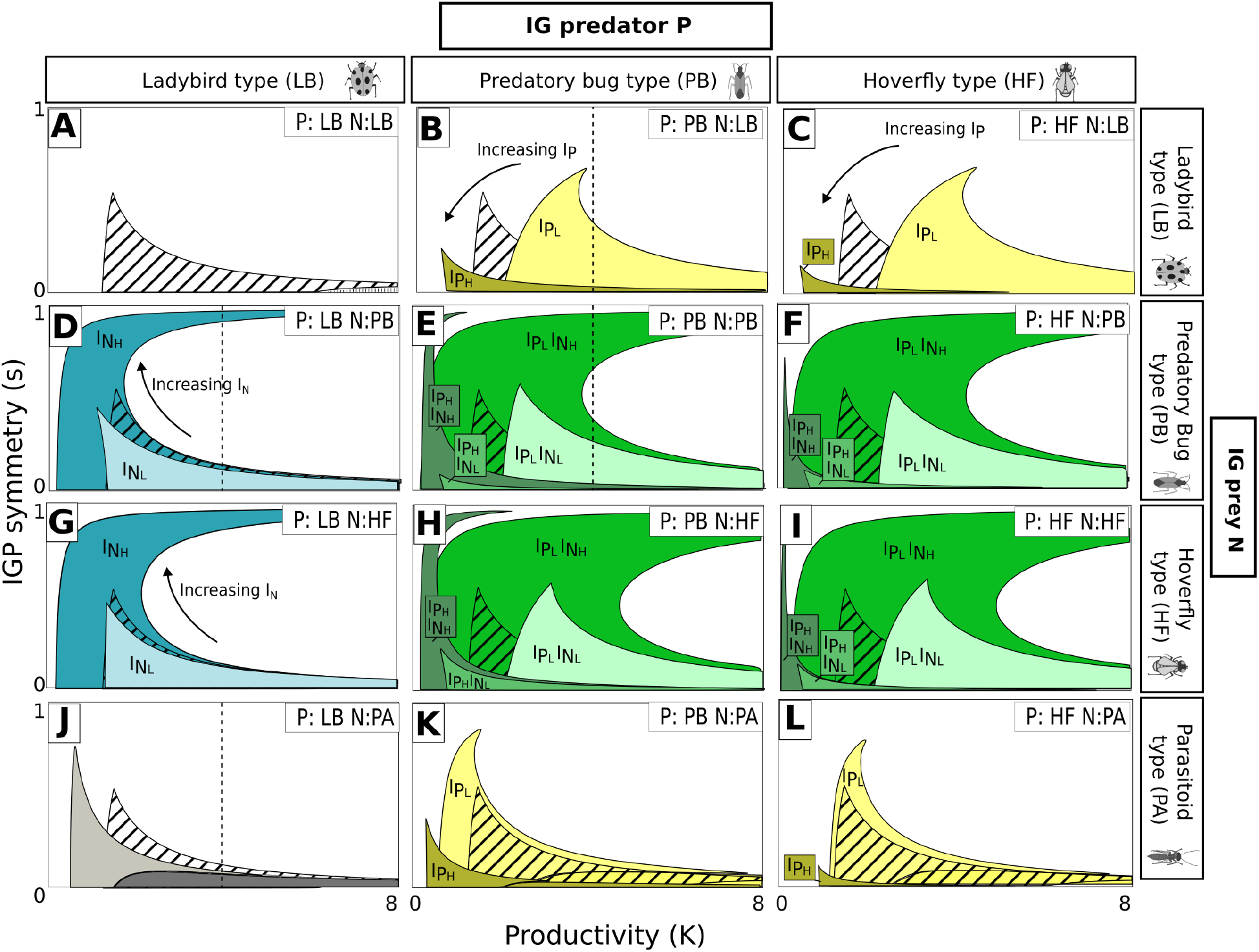
Changes in coexistence across IGP modules with different predator types. Coexistence (*HNP* * equilibria) region along IGP symmetry *s* and productivity *K* for all 12 combinations of IG predators (LB, PB and HF) and IG preys (LB, PB, HF, PA) and two levels of external input rates (low and high). We used colors to group the changes in comparison to **LBLB** baseline module (**A**): when the IG predator *P* was changed to a PB or HF type (**B, C, K, L**, yellow colors), when the IG prey *N* was a PB or HF type (**D, G**, blue colors), and when both IG predator *P* and IG prey *N* belong to the PB and HF types (**E, F, H, I**, green colors). The gradient within each color represents the combination of high (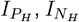) and low external input rates (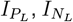) for both the IG predator *P* and the IG prey *N*. Low and high values of *I*_*P*_ and *I*_*N*_ were 10 and 90 % percentile of the maximum input for each predator type (Table 1). In the **LBPA** module (**J**), light and dark gray represent stable and periodic coexistence areas, respectively. For each plot we added the **LBLB** coexistence area as a baseline (hashed area). Dotted vertical lines represent transects shown in figure 4, for fixed high productivity (*K* = 4).

**Figure 4:**
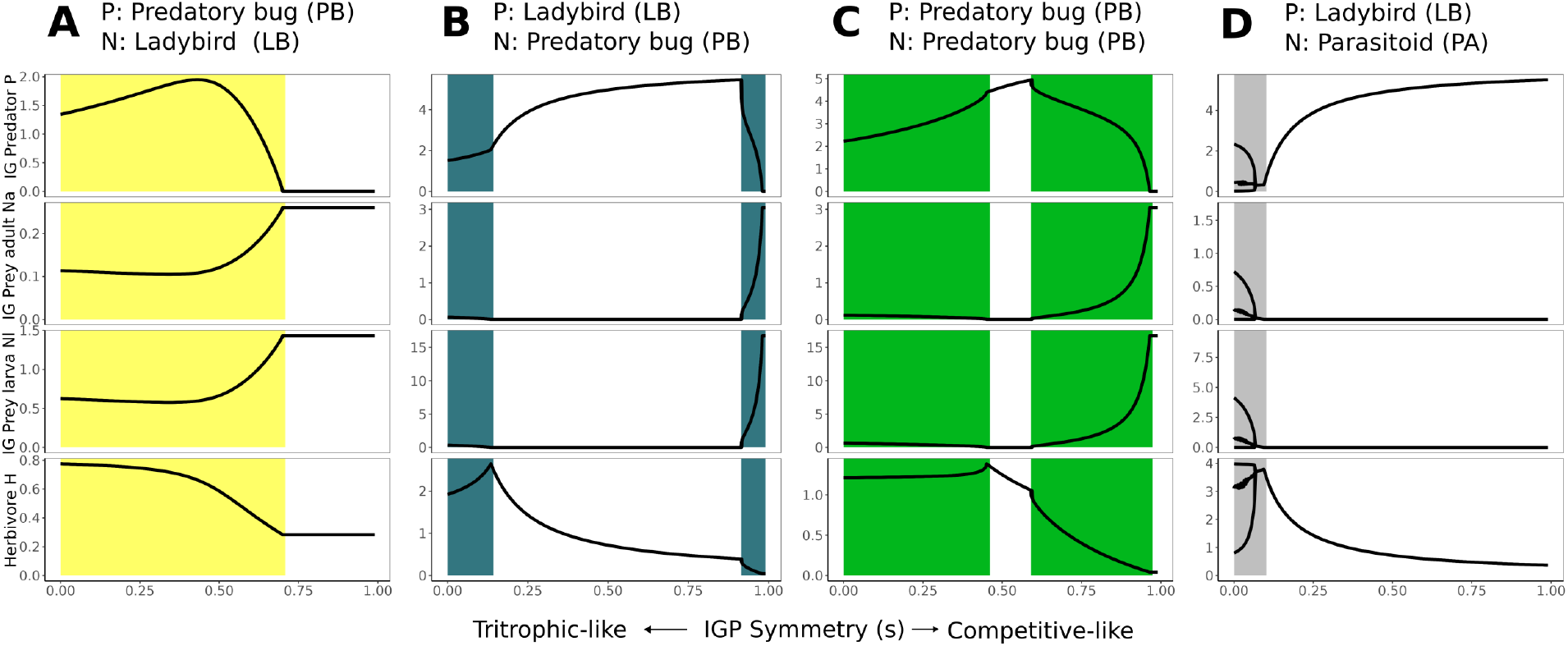
Equilibrium population densities of four IGP modules along the IGP symmetry gradient *s*. We show the population densities at equilibria of the IG predator *P*, IG prey adult *N*_*a*_ and larva *N*_*l*_ and the herbivore *H* for the **PBLB** (**A**), the **LBPB** (**B**), the **PBPB** (**C**) and the **LBPA** (**D**) modules along the IGP symmetry *s* gradient. For all modules, we have *K* = 4, 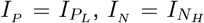. Colored regions represent the regions of coexistence (*HNP* * equilibrium) and follow the same color code as in f figure 3. In the **LBPA** module, we represent only the maximum and the minium value of the periodic equilibrium and the stable point (as we have a bistability of a periodic and fixed point).

#### 3.2.1 Predatory Bug and Hoverfly as IG predator and IG prey types

When we included the Predatory Bug (PB) or the Hoverfly (HF) types as IG predators *P*, changes in the coexistence region depended on the external input rate *I* (modules **PBLB** and **HFLB**, yellow colors in fig.3B and C). Low quantities of external input for the IG predator 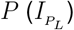 increased the area of the region of coexistence in relation to the **LBLB** scenario and shifted it to higher productivities. This means that when IG predators *P* of PB or HF types required high productivities *K* to invade the IG prey *N* (left boundary of the coexistence area), they would need even a higher *K* to exclude the IG prey *N* (right boundary of the coexistence area). Within the enhanced coexistence area, species equilibria varied non-linearly with *s* (Fig. 4A). At low-to-medium values of *s* (0 to 0.5, tritrophic-like to symmetric IGP modules) the herbivore declined slightly as the IG predator *P* increased, while the IG prey *N* decreased slowly. When the IGP module became more competitive than symmetric (*s >* 0.5), *N* was released, overcoming predation and out-competing *P*. This caused an abrupt decline in the herbivore *H* and consequently the IG predator *P* persisted via the external input *I*_*P*_ contrarily to the **LBLB** module. When we increased the external input 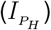, the IG predator *P* invaded and excluded the IG prey *N* at lower productivities than in the **LBLB** module. Here, the coexistence region collapsed and was only possible with very low values of *s* (almost tritrophic chains), where *P* relied heavily on the *N* (Fig.3 B and C)).

In the IGP modules where we included the PB and HF as IG prey *N*, coexistence also depended on the external input *I*_N_. With low values of external input (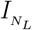), the area of coexistence was not really affected. But, as soon as we increased it (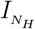), the coexistence area expanded significantly (modules **LBPB** and **LBHF** in Fig.3D and G, blue colors). The left limit of the coexistence region shifted to the left: this implied that the IG predator *P* could invade the system where the IG prey *N* was present at lower values of productivity, and even for high values of IGP symmetry (*s >* 0.6). For high values of IGP symmetry (this is, for competitive-like modules with low intraguild predation), coexistence was possible for a large range of productivity. This region of coexistence was not present in the **LBLB** module. The dependence of the population equilibria on *s* was similar to the **LBLB** module at near-tritrophic-like IGP (*s ≈* 0) 4B vs fig. 2C): herbivore *H* and IG predator *P* densities increased with *s*, reducing the IG prey *N*. However, in the coexistence region for near-competitive networks (*s* → 1), this trend reversed. The external input *I*_*N*_ enabled the coexistence of both predators even in a competitive scenario with low intraguild predation. As *s* increased, the system became more competitive allowing the superior competitor (IG prey *N*) to increase while the IG predator *P* decreased, ultimately reducing the shared herbivore *H* density (Fig. 4B).

When the predator types PB or HF acted both as IG predator *P* and IG prey *N*, the result was a combination of the previous cases (modules **PBPB, HFPB, PBHF** and **HFHF** in Fig.3E, F, H and I, green colors). When both external inputs *I*_*N*_ and *I*_*P*_ were low (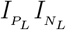), there was no effect on coexistence, except for a shift to higher productivity values. If the input *I*_*P*_ for the IG predator *P* was low, whereas the input *I*_N_ for the IG prey *N* was high (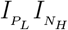), coexistence was highly enhanced, even for symmetric IGP (*s* → 0.5). However, when the external input rate to the IG predator *I*_*P*_ was higher (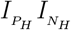 and 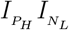), coexistence was limited to low productivities, or to highly tritrophic-like IGP modules. Within the region of coexistence in the best case scenario (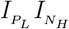), *P* density increased monotonically while *N* slowly decreased with increasing *s* (Fig. 4C). Notably, herbivore density remained relatively constant, as IG predator *P* ‘s slow increase was compensated by the IG prey *N* ‘s decline. In more competitive-like networks (*s >* 0.5), the combination of external inputs drove an increase in *N* and a decrease in both *P* and *H*.

#### 3.2.2 Parasitoid as IG prey type

When the IG prey *N* belonged to the parasitoid (PA) type with the LB type as IG predator, the coexistence region shifted to lower values of productivity, and to slightly higher values of IGP symmetry, compared to the baseline **LBLB** module (Fig.3J, K, L, grey colors). The periodic coexistence region increased and shifted to lower values of *K*. When PB and HF types were included as IG predators, we got similar effects as the ones described for the **PBLB** and the **HFLB** modules. Low and medium values of *I*_*P*_ increased the coexistence for low and medium productivities, and high values of *I*_*P*_ collapsed the region of coexistence (Fig.3K and L, yellow colors). For near tritrophic-like modules (*s* → 1), the system had bistability equilibria between the fixed point and the periodic coexistence solutions. The amplitude of the oscillation for each variable slowly decreased with increasing *s*, collapsing at higher herbivore densities (Fig. 4D).

### 3.3 Herbivore suppression across IGP modules

So far, we have explored the role of different predator types and IGP symmetry on the coexistence of predators along a productivity gradient. Our aim now is to compare the relative efficacy of herbivore suppression within and outside coexistence across the 12 studied IGP modules. For each IGP module, we extracted the population density of the herbivore along the productivity gradient *K* for three values of IGP symmetries *s* that represented tritrophic-like (*s* = 0.1), symmetric (*s* = 0.5), and competitive-like (*s* = 0.9) modules. We also considered population densities for external input *I*_*x*_ that maximized the coexistence region for the Predatory bug and the Hoverfly types as predators (this is, a low *I*_*P*_ and a high *I*_*N*_, see Fig. 3). We first describe the role of the IGP symmetry in herbivore suppression and then the influence of different predator types, in comparison to the baseline **LBLB** module.

#### 3.3.1 IGP symmetry and herbivore suppression

In all IGP modules, herbivore suppression along productivity was mainly driven by the IGP symmetry (Fig. 5). For symmetric and competitive-like modules (*s* = 0.5 and *s* = 0.9; with lower intraguild predation and higher competition), the herbivore density was kept low across an increasing gradient of productivity (circles and squares shapes in Fig. 5). This was not significantly affected by the coexistence of both predators (filled shapes in Fig. 5D, E, F, G, H and I). In these scenarios, the IG prey *N* has higher and slowly decreasing values with increasing productivity *K* (section 4 in the Supplementary Material), keeping the herbivore densities low. For tritrophic- like modules (with high intraguild predation and low competition, *s* = 0.1), the herbivore density increased considerably within the region of coexistence (filled shapes in all panels of Fig. 5) and stabilized at high values outside the region of coexistence as *K* increased (empty shapes in Fig. 5 A, D, G). The stabilization was visible only if coexistence stopped within the explored region of productivity (*K* ≤ 8). Within the coexistence region, the increase of the herbivore was led by the low and decreasing values of the IG prey *N* (section 4 in the Supplementary Material).

**Figure 5:**
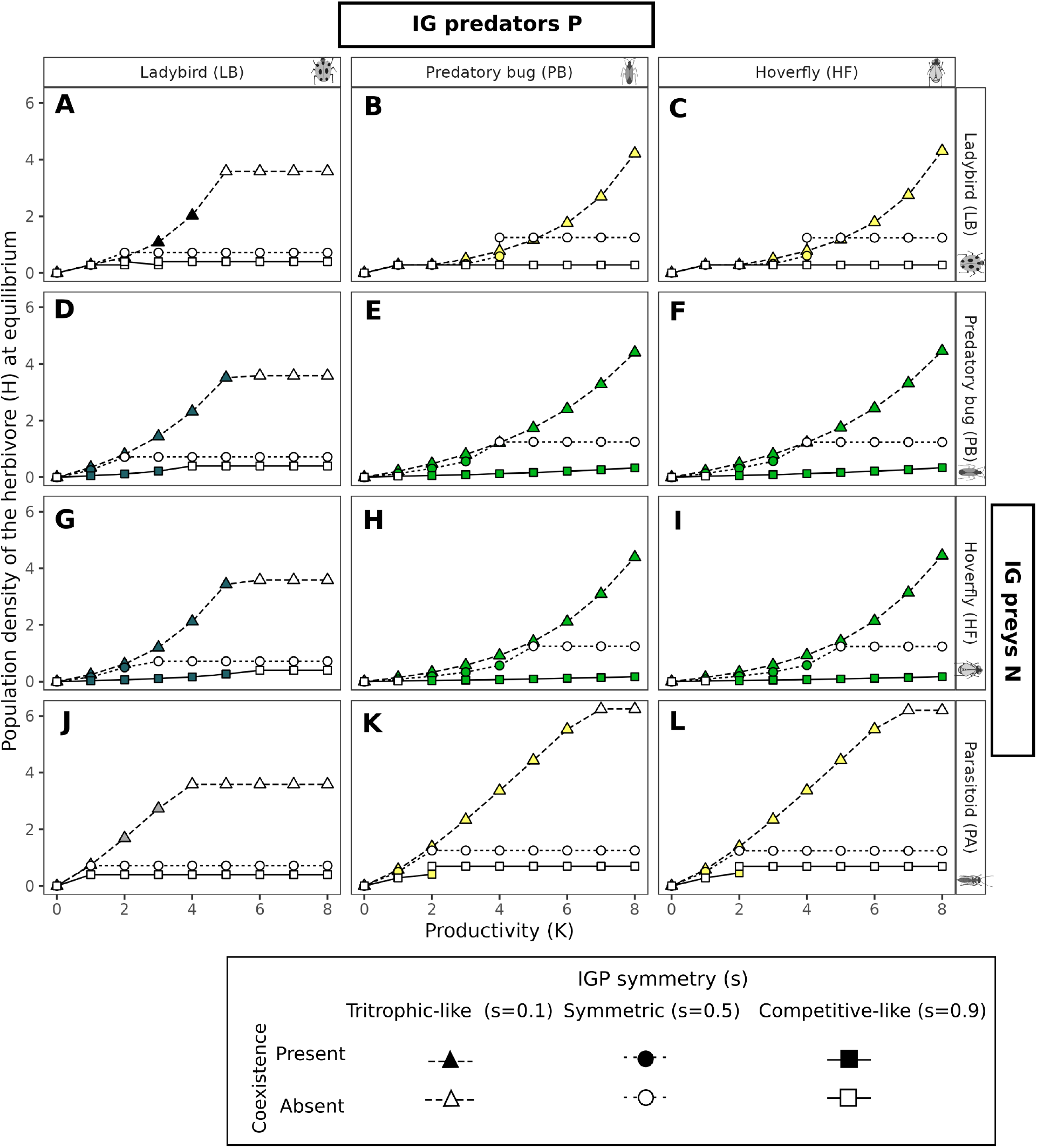
Herbivore suppression and natural enemies’ coexistence. We show the population density of the herbivore (*H*) at equilibrium for each of the 12 IGP modules, along the productivity gradient and for three representative IGP symmetries: tritrophic-like (*s* = 0.1, triangles and dashed lines), symmetric (*s* = 0.5, circles and short-dashed line) and competitive-like (*s* = 0.9, squares and continuous line). We represent if the system was at a coexistence equilibria (*HNP* ^*^ equilibrium, filled shapes) or not (without distinguish between *H*^*^, *HN* ^*^ or *HP* ^*^ equilibria, empty shapes). For the IGP modules that include the predatory bug (PB) or the hoverfly (HF) types as IG predators or IG prey (**B, C, D, G, E, H, F, I, K** and **L**), we include only the values of external input 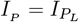 and 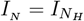 that increased coexistence. Colors of the filled shapes follow the color code of figure 4.

#### 3.3.2 Different predator types, coexistence and herbivore suppression

The area of the coexistence region across different IGP symmetries and productivities, and its subsequent effect on herbivore suppression, depended on the predator types used. In the **LBLB** baseline module, minimal herbivore densities occurred in competitive-like and symmetric modules outside the region of coexistence where only one predator was present (empty circles and squares in Fig. 5A). Within the region of coexistence of tritrophic-like modules, we observed the release of the herbivore with increasing productivity (filled triangles in Fig. 5A). Including HF or PB as IG predators *P* with a low external input *I*_*P*_ shifted the coexistence region to higher values of *K* in the tritrophic-like modules (filled triangles in Fig. 5B and C). This resulted in a lower herbivore density in comparison to the **LBLB** module, for a certain range of *K*. However, at very high productivity *K*, this lead to higher values of herbivore density. Thus, the highest herbivore suppression across increasing productivities was still found outside coexistence and for competitive-like modules (empty circles and squares, Fig. 3B and C). When PB and HF acted as IG prey *N* (**LBPB** and **LBHF**) with a high external input *I*_*N*_, coexistence was extended to higher productivities compared to the **LBLB** baseline, both for tritrophic and competitive-like modules (filled triangles and squares in Fig. 5D and G). For tritrophic-like modules, this resulted again in high herbivore densities due to predator release. In competitive-like modules, predator release was not significant within coexistence as the IG prey *N* was kept low (section 4 in the Supplementary Material). In this sense, coexistence of predators and minimal values of herbivore density were found over a wide range of productivity, especially in the **LBHF** module (Fig. 5G). In modules where PB (or HF) acted *both* as IG predator *P* and IG prey *N* (with low *I*_*P*_ and high *I*_*N*_), we observed the highest herbivore suppression in competitive-like modules, where predators coexistence was greatly enhanced (filled squares in Fig. 5E, F, H and I). For tritrophic-like modules, an increase in the herbivore density was again found (filled triangles). Finally, when the Parasitoid type acted as the IG prey *N* (regardless of IG predator, **LB, PB** or **HF**), the herbivore density within the coexistence region was consistently higher than in the **LBLB** baseline and increased dramatically with productivity ((Fig. 5J, K and L)). Here, the minimal value of herbivore density occurred outside the coexistence region.

## 4 DISCUSSION

Coexistence of multiple predators in animal communities and their combined effects on the suppression of shared prey are usually difficult to predict (Chang and Cardinale, 2020; Brodeur and Boivin, 2006; McCoy et al., 2012), given that predators are embedded in complex networks of competitive, mutualistic and predatory interactions (Chailleux et al., 2014) where dynamics depart from predictions based on pairwise expectations (McCoy et al., 2012). Even in the specific case of intraguild predation, predictions from relatively simple IGP models of two predators that prey on the same species, but also feed on each other, are not always in line with empirical observations of IGP modules. Here, we tested the hypothesis that greater functional dissimilarities between intraguild predators could overcome the interference created by intraguild predation and exploitative competition, enhancing their coexistence as well as their effect on prey suppression (Snyder, 2019; Alhadidi et al., 2019). We modeled this in an empirically inspired setup to identify how such functional differences between natural enemies (predators and parasitoids) affect their coexistence conditions and modify herbivore suppression. We found that functional differences between natural enemies enhance coexistence across different IGP symmetries and productivity levels, especially when these predators can feed on external resources. In competitive-like IGP modules (where the IG predator feeds mostly on the shared prey), this leads to scenarios with the strongest herbivore suppression, even when both predators are present.

### 4.1 Coexistence depends on IGP symmetry and productivity

For all 12 IGP modules composed of pairs of functionally different predator types, coexistence of natural enemies along a productivity gradient *K* depends on IGP symmetry *s*. Even though the influence of IGP symmetry and productivity has been partially analyzed and described before (Sentis et al., 2013; Vandermeer, 2006; Hin et al., 2011; Diehl and Feißel, 2000; Mylius et al., 2001; Sentis et al., 2014), our study draws a complete picture of their interaction. In the **LBLB** baseline module, composed of ladybirds without prey-structure nor alternative prey, the region of coexistence of predators is restricted from perfectly symmetrical to tritrophic-like modules (*s <* 0.5) as productivity increases. This result confirms previous findings on the limitation of coexistence to intermediate levels of productivity for stage-structured models (Diehl and Feißel, 2000; Hin et al., 2011; Mylius et al., 2001) and reaffirms that IGP symmetry, especially tritrophic- like modules, can stabilize coexistence in enriched IGP systems (Faria and Costa, 2009). In our model, the region of coexistence is also a region of mutual invasibility, where both predators can invade a system where the other predator is present. This implies that, given a specific IGP symmetry *s*, coexistence is possible only if the minimum value of productivity *K* needed for the IG predator *P* to invade a *HN* system is lower than the maximum value of *K* above which the IG prey *N* can not longer invade a *HP* system (see SM 1 for the mathematical derivation and Diehl and Feißel (2000) for equivalent results). This hypothesis could be experimentally verified in settings where productivity *K* is manipulated (as in (Diehl and Feißel, 2000)), or in real communities across productivity gradients as in (Novak, 2013).

The inclusion of predators from the Predatory Bug (PB) and Hoverfly (HF) types with an external food input modifies the coexistence regions compared to the **LBLB** baseline module. Consistent with previous work, an external input to the IG predator (*I*_*P*_) generally hinders coexistence, while an input to the IG prey (*I*_*N*_) enhances it (Holt and Huxel, 2007; Daugherty et al., 2007; Briggs and Borer, 2005; Van Rijn et al., 2002). However, in our system, we assumed that, both in PB and HF types, half of food intake during their life-cycle relies on this input. As a result, the effect of external input on the IG predator and IG prey was mediated by its average rate. A low input rate for the IG predator (*I*_*P*_) expands coexistence to a broader and higher range of productivity compared to the baseline **LBLB** module, although coexistence remains restricted to tritrophic-like modules (where the IG predator feeds primarily on the IG prey). In contrast, a high input for IG prey *I*_*N*_ allows coexistence for competition-like modules (*s >* 0.5, where the IG predator feeds primarily on the shared herbivore and intraguild predation is low) for a wide range of productivity, which is not possible for the **LBLB** module. In this sense, external food sources may relax either the trophic cascade triggered by intraguild predators in nearly tritrophic systems or allow poor competitors to persist in purely competitive interaction modules. The external input for the IG prey *N* allows the IG predator *P* to invade at lower values of productivity *K* for any given symmetry *s*, thus, increasing the area of coexistence. These results are relevant for productive agricultural systems (Amarasekare, 2008) and show how supplementing external food sources can stabilize coexistence (Shchekinova et al., 2013; Van Rijn et al., 2002; Briggs and Borer, 2005). A recent meta-analysis has explored how external sources like pollen, infertile insect eggs (both commonly used to feed natural enemies in the context of biocontrol), or alternative prey usually increase predator populations and ultimately benefit herbivore suppression by single natural enemies (Deere et al., 2024). Other studies on predatory mites have shown that pollen reduces intraguild predation in assemblages of multiple predators, often resulting in longer predator coexistence (Tsuchida et al., 2022), although null effects on coexistence have also been reported (Abad-Moyano et al., 2010).

Interestingly, a Parasitoid (PA) predator type does not enhance coexistence in comparison to the **LBLB** baseline module. Consistent with earlier findings, parasitoids as IG prey shift coexistence towards lower productivity (Borer et al., 2007) and increase instability at high productivity levels (Nakazawa and Yamamura, 2006). This result is related to the lack of additional mechanisms that would allow the parasitoid (the intraguild prey) to overcome competition and predation by the intraguild predator. Its functional differences do not result in any relaxation of competition; instead, they increase coincidental intraguild predation. Using an experimental and theoretical framework, it was shown that this outcome can be reversed, if the intraguild prey (a parasitoid) has temporal refuges that enable it to survive at higher productivities (Amarasekare, 2008, 2007).

### 4.2 Coexistence and functional differences can lead to strong herbivore suppression

The impact of coexistence of natural enemies on herbivore suppression was strongly modulated by IGP symmetry. For tritrophic-like modules (with high intraguild predation and low competition), herbivore suppression was strong at low values of productivity even within the region of coexistence (“High herb. supp.” and green cells in Table 3). However, we observed that an IG predator-release mechanism within coexistence leads to a negative impact of intraguild predation on herbivore suppression as shown before (Rosenheim and Harmon, 2006; Diehl and Feißel, 2000; Vance-Chalcraft et al., 2007). This leads to high herbivore densities at high productivities either within coexistence, or where only the inferior competitor- the IG predator *P* - is present (“Low. herb. supp.” in Table 3). In accordance to these results, in a system of thrips with two predator mites with high intraguild predation (*Neoseiulus cucumeris* and *Amblyseius swirskii*), both predators persisted for up to four weeks, but their combined effect did not improve thrips suppression (Delisle et al., 2015). On the contrary, we found the highest herbivore suppression in symmetric (*s* = 0.5) and competitive-like modules (*s* = 0.9) across all productivities, regardless of coexistence (Table 3). This makes sense when coexistence is not possible, as the IG prey *N* excludes the IG predator *P* and drives herbivore density to its minimum being the superior predator (Mylius et al., 2001; Hin et al., 2011; Diehl and Feißel, 2000). However, herbivore densities are also kept low when coexistence is possible in some of the IGP modules with different predator types (Table 3). In the **LBLB** baseline module, one of the two predators excludes the other depending on IGP symmetry, on the increase of productivity- which reduces the impact of competition between both predators-, and even on the initial conditions (in the bistability region). However, when PB or HF act as IG prey in symmetric and competitive-like modules, the external input to the IG prey balances the predation pressure from the IG predator and at the same time reduces the pressure on the shared herbivore. This results in high IG prey densities and strong herbivore suppression within the coexistence region. This effect is even stronger when PB or HF act as IG predators with a low IG predator input: in these scenarios co- existence and herbivore suppression are extended over a broader productivity range (Table 3).In short, our model shows strong herbivore suppression upon IG predator addition in competitive-like modules because: (1) the intraguild predation is relatively low, and (2) even if the structural competition is high, the external food sources reduce competition between both natural enemies, keeping their population densities higher. Therefore, as other authors have pointed out, adding generalist or stage-structured predators can stabilize and make coexistence possible for highly competitive IGP systems (with a relatively low intraguild predation) while providing a strong herbivore suppression (i.e effective long-term biocontrol services, Faria et al. (2008)). In cotton, a long-term study with generalist predatory bugs *Geocoris spp*. and *Orius tristicolor* found that both predators reduced the abundance of a specialist predatory mite *Galendromus occidentalis* (here the IG prey), yet improved overall spider mite suppression (Colfer et al., 2003). The pattern is also consistent with greenhouse evidence showing that coexistence with effective herbivore suppression can emerge at low IGP levels with broader resource availability, where the species *Macrolophus pygmaeus* (IG prey) and *Orius laevigatus* (IG predator) both capable of feeding on plant resources, coexisted for months and together suppressed thrips and aphids more effectively than either predator alone (Messelink and Janssen, 2014).

**Table 3:** Overview of the effect of different predator types on coexistence and herbivore suppression across IGP modules. We summarize how IGP combinations of different predator types affect the coexistence of predators and the suppression of the herbivore along low and high productivities (*K* ∈ [1 − 4] and *K* ∈ [5 − 8]), as well as along three IGP symmetries *s* (tritrophic (*s* = 0.1), symmetric (*s* = 0.5) and competitive-like (*s* = 0.9)). We grouped the predatory bug PB and hoverfly HF types combinations for the external inputs rates scenarios that maximized coexistence (low *I*_*P*_ and high *I*_*N*_). For each IGP module, we annotate if the coexistent was present (“Coexistence”) or absent (“No coexistence”) in more than half of the range. Herbivore suppression was high (“High herb. supp.”) or low (“Low herb. supp.”) if the mean herbivore density was lower or higher than 1.8 (which represents half of the maximum value of the herbivore density obtained in the **LBLB** baseline) (see Supplementary table S3). Within each group, the colored cells highlight where coexistence was present with a low (yellow) or a high (green) herbivore suppression. (*) Coexistence was not present in this combination when the IG Predator was from the LB type

| | IG predator P | IG prey N | IGP symmetry ($s$) and Productivity ($K$) | | | | | |
| --- | --- | --- | --- | --- | --- | --- | --- | --- |
|  |  |  | Tritrophic-like module |  | Symmetric module |  | Competitive-like module |  |
|  |  |  | Low Product. | High Product. | Low Product. | High Product. | Low Product. | High Product. |
| IGP modules | Ladybird **LB** | Ladybird **LB** | Coexistence<br>High herb. supp. | No coexistence<br>Low herb. supp. | No coexistence<br>High herb. supp. | No coexistence<br>High herb. supp. | No coexistence<br>High herb. supp. | No coexistence<br>High herb. supp. |
|  | Predatory Bug **PB**<br>Hoverfly **HF** | Ladybird **LB** | No coexistence<br>High herb. supp. | Coexistence<br>Low herb. supp. | No coexistence<br>High herb. supp. | No coexistence<br>High herb. supp. | No coexistence<br>High herb. supp. | No coexistence<br>High herb. supp. |
|  | Ladybird **LB** | Predatory Bug **PB**<br>Hoverfly **HF** | Coexistence<br>High herb. supp. | No coexistence<br>Low herb. supp. | Coexistence<br>High herb. supp. | No coexistence<br>High herb. supp. | Coexistence<br>High herb. supp. | No coexistence<br>High herb. supp. |
|  | Predatory Bug **PB**<br>Hoverfly **HF** | Predatory Bug **PB**<br>Hoverfly **HF** | Coexistence<br>High herb. supp. | Coexistence<br>Low herb. supp. | Coexistence<br>High herb. supp. | No coexistence<br>High herb. supp. | Coexistence<br>High herb. supp. | Coexistence<br>High herb. supp. |
|  | Ladybird **LB**<br>Predatory Bug **PB**<br>Hoverfly **HF** | Parasitoid **PA** | Coexistence<br>Low herb. supp. | Coexistence (*)<br>Low herb. supp. | No coexistence<br>High herb. supp. | No coexistence<br>High herb. supp. | No coexistence<br>High herb. supp. | No coexistence<br>High herb. supp. |

When PA acted as the IG prey, herbivore suppression is low when coexistence is possible. This occurs in tritrophic-like modules, where the IG predator mostly feeds on the larval stage of the IG prey leading to a negative effect on adult PA that reduces the control of the herbivore (Table 3, and as recently found by Martínez-Martínez et al. (2026)). Instead, high herbivore suppression is found outside coexistence in symmetric and competitive-like modules with parasitoids, when only one predator is present. These results match previous theoretical findings (Nakazawa and Yamamura, 2006), but contradict the predictions provided by more recent models in which herbivore suppression is higher when both natural enemies are present (Alves-Rubio et al., 2022), probably due to the fact that these models consider predators with three ontogenetic stages (eggs, larva and adult). Yet, a compelling experimental setup (Alhadidi et al., 2019) has shown that combining predators and parasitoids does not necessarily enhance herbivore suppression, in agreement with our model predictions.

### 4.3 Functional types, coexistence and herbivore suppression in IGP

Our findings show that combining different predator types modify coexistence and herbivore suppression compared to the same predator type **LBLB** baseline (Table 3). This should confirm the dissimilarity hypothesis, where functional differences between predators increase their complementarity, dilute antagonistic interactions, and ultimately promote coexistence (Schmitz, 2007; Snyder, 2019; Diehl et al., 2013). However, we also find that the effects of functional differences are nuanced by: i. the presence of an external input, ii. the position of the natural enemy in the IGP module (IG predator or as an IG prey), iii. productivity, and iv. IGP symmetry. Notably, the best scenarios for enhanced coexistence and herbivore suppression occurs when the HF and PB types acted as both IG prey and IG predator, particularly for competitive-like modules (Table 3). However, this also happens in modules with the same predator type (**PBPB** and **HFHF**), in contradiction with the dissimilarity hypothesis, but the parameter space for coexistence and herbivore suppression in the same predator type scenarios seems to be unrealistic. On the contrary, for IGP modules composed of functionally dissimilar predators (**PBHF** and **HFPB**), parameter conditions for coexistence and herbivore suppression are more plausible.

Experimental work has both validated or contradicted our results. For example, studies combining PB and HF predator types showed that when *Orius majusculus* and *O. laevigatus* (PB types) acted as IG predators, they reduced the populations of the specialist midge *Aphidoletes aphidimyza* (HF type). Although coexistence of the predators was compromised, aphid suppression was not reduced (Messelink and Janssen, 2014). By contrast, when a predatory bug (PB) acts as the IG prey in combination with a ladybird (LB), persistence is less likely, as shown in the *Macrolophus pygmaeus–Adalia bipunctata* system, where the predatory bug could no longer persist once the shared prey *Acyrthosiphon pisum* declined (Trotta et al., 2015). These discrepancies reveal the importance of experimental designs suited to test the hypotheses numerically explored here, for example by testing under similar conditions various guild combinations.

### 4.4 Final comments

Building a model based on commonly used empirical natural-enemy aphid system helped us explore the mechanisms underlying the role of functional different predator types on coexistence and shared prey suppression (Alhadidi et al., 2019; Snyder, 2019). Clearly, our simplified three species IGP modules should not allow for precise predictions as they depend on the way we incorporated interactions and parameters Vandermeer (2006). Our models also ignore reciprocal intraguild predation (where both predators can feed on the other, (Rosenheim et al., 1995)). They highlight, though, how functional differences between predators interact with IGP symmetry and productivity leading to different coexistence and herbivore suppression outcomes. Our analysis also ignored transient dynamics or unstable equilibria (e.g. cycles), which are particularly relevant for agricultural systems that are constantly subjected to seasonal changes, demographic stochasticity or harvesting perturbations (Rosenheim and Harmon, 2006; Briggs and Borer, 2005; Frago and Godfray, 2014). This can also change the relative role of predator types, as generalist predator types are more prone to survive in changing agricultural systems (Diehl et al., 2013). More importantly, the predator types we used might be insufficient to capture all functional differences among natural enemies, ignoring for instance hunting strategies, prey preference or spatial movement (Alhadidi et al., 2019; Krivan and Diehl, 2005). Despite focusing on the aphid system, our predator types still capture much of the observed structures in arthropod IGP modules, and can therefore be extrapolated to other systems of herbivores and their natural enemy predators. The different patterns we find in the simple ecological modules we study worth to be compared with approaches that focus on the whole community of natural enemies (Jonsson et al., 2017), with meta-analysis of previous studies (Diehl et al., 2013), as well as with lab microcosm experiments (Briggs and Borer, 2005), and controlled pilot implementations in the field. Only at the crossing of these different approaches, can we gain a deeper understanding of ecological processes that maintain coexistence in multi-predator communities while enhancing long-term biocontrol services in agroecological systems.

## ACKNOWLEDGMENTS

This work was funded by the ANR PRCE project (22-CE32-0009): ENEMYCOCKTAIL: Designing natural enemy cocktails for a better biocontrol. EMVC thanks all the members of BioDICee lab (and beyond) for the comments and ideas, the faries and H. Roland for the help and support during the whole process.

## AUTHOR CONTRIBUTIONS

Conceptualization: EMVC, VD, EF and FM. Methodology: EMVC and VD. Investigation: EMVC. Writing-–original draft: EMVC and VD. Writing-–review & editing: VD, EF and FM. Funding acquisition: EF and VD. Resources and supervision: VD.

