## Supplementary material for "Predator coexistence and herbivore suppression are shaped by predator functional types in intraguild predation modules"

#### 1 Additional methods: IGP modules equations and structures

In this section, we show the equations for all IGP modules. We write down the **LBLB** baseline and then the result of changing the IG predator P (red color) and the IG prey N (blue color). See figure S.3 for all the visual representations.

##### 1.1 Baseline

###### 1.1.1 LBLB (IG Predator P = Ladybird LB, IG prey N = Ladybird LB)

$$
\begin{aligned}
\frac{dP}{dt} &= e_{P}(f_{NP(H,N_l)} + f_{HP(H,N_l)})P - \mu_{P} P \\
\frac{dN_a}{dt} &= m_{N} N_l - \mu_{N} N_a \\
\frac{dN_l}{dt} &= e_{N} f_{HN(H)}(N_l + N_a) - f_{NP(H,N_l)}P - (\mu_{N} + m_{N})N_l \\
\frac{dH}{dt} &= \rho(K - H) - f_{HN(H)}(N_l + N_a) - f_{HP(H,N_l)}P
\end{aligned}
$$

Where $P$ represents the IG predator, $N_a$ and $N_l$ represent the adult and larval stages of the IG prey and $H$ the herbivore. $e_P$ and $e_N$ are the conversion efficiencies, $\mu_P$ and $\mu_N$ the natural death rates and $m_N$ the maturation rate of $P$ and $N$, $s$ is IGP symmetry (relative feeding on H or N), $\rho$ the chemostat turnover and $K$ the productivity of the system. $f_{XY}$ are the functional responses of $Y$ feeding on $X$, where $Y \in \{P, N\}$ and $X \in \{N, H\}$. Their formulation is at the end of the section.

#### 1.2 Changing the IG predator P

##### 1.2.1 PBLB (IG Predator P = Predatory bug PB, IG prey N = Ladybird LB)

When the IG predator P belongs to the predatory bug type, we add an external input rate $I_P$ with preference $\phi_P$ to the IG predator. (where $\phi_P = 0.5$).

$$
\begin{aligned}
\frac{dP}{dt} &= e_P((1-\phi_P)(f_{NP(H,N_l)} + f_{HP(H,N_l)}) + \phi_P I_P)P - \mu_P P \\
\frac{dN_a}{dt} &= m_N N_l - \mu_N N_a \\
\frac{dN_l}{dt} &= e_N f_{HN(H)}(N_l + N_a) - (1-\phi_P) f_{NP(H,N_l)} P - (\mu_N + m_N) N_l \\
\frac{dH}{dt} &= \rho(K - H) - f_{HN(H)}(N_l + N_a) - (1-\phi_P) f_{HP(H,N_l)} P
\end{aligned}
$$

##### 1.2.2 HFLB (IG Predator P = Hoverfly HF, IG prey N = Ladybird LB)

When the IG predator P belongs to the hoverfly type, we add an stage structure for the IG predator P where only the larva $P_l$ fed on the aphid and the adult $P_a$ fed on an external input $I_P$

$$
\begin{aligned}
\frac{dP_a}{dt} &= m_P P_l - \mu_P P_a \\
\frac{dP_l}{dt} &= e_P((f_{NP(H,N_l)} + f_{HP(H,N_l)})P_l + I_P P_a) - (\mu_P + m_P) P_l \\
\frac{dN_a}{dt} &= m_N N_l - \mu_N N_a \\
\frac{dN_l}{dt} &= e_N f_{HN(H)}(N_l + N_a) - f_{NP(H,N_l)} P_l - (\mu_N + m_N) N_l \\
\frac{dH}{dt} &= \rho(K - H) - f_{HN(H)}(N_l + N_a) - f_{HP(H,N_l)} P_l
\end{aligned}
$$

#### 1.3 Changing the IG prey N

##### 1.3.1 LBPB (IG Predator P = Ladybird LB, IG prey N = Predatory Bug PB)

When the IG prey N belongs to the predatory bug type, we add an external input rate $I_N$ with preference $\phi_N$ to the IG prey (where $\phi_N = 0.5$)..

$$
\begin{aligned}
\frac{dP}{dt} &= e_P(f_{NP(H,N_l)} + f_{HP(H,N_l)})P - \mu_P P \\
\frac{dN_a}{dt} &= m_N N_l - \mu_N N_a \\
\frac{dN_l}{dt} &= e_N((1-\phi_N) f_{HN(H)} + \phi_N I_N (N_l + N_a)) - f_{NP(H,N_l)} P - (\mu_N + m_N) N_l \\
\frac{dH}{dt} &= \rho(K - H) - (1-\phi_N) f_{HN(H)}(N_l + N_a) - f_{HP(H,N_l)} P
\end{aligned}
$$

##### 1.3.2 LBHF (IG Predator P = Ladybird LB, IG prey N = Hoverfly HF)

When the IG predator N belongs to the hoverfly type, only the larva $N_l$ fed on the aphid and the adult $N_a$ fed on an external input $I_P$

$$
\begin{aligned}
\frac{dP}{dt} &= e_P(f_{NP(H,N_l)} + f_{HP(H,N_l)})P - \mu_P P \\
\frac{dN_a}{dt} &= m_N N_l - \mu_N N_a \\
\frac{dN_l}{dt} &= e_N(f_{(HN(H)}N_l + I_N N_a) - f_{NP(H,N_l)}P - (\mu_N + m_N)N_l \\
\frac{dH}{dt} &= \rho(K-H) - f_{HN(H)}N_l - f_{HP(H,N_l)}P
\end{aligned}
$$

#### 1.4 Changing both (summary of previous cases) between PB and HF

##### 1.4.1 PBPB (IG Predator P = Predatory bug PB, IG prey N = Predatory bug PB)

$$
\begin{aligned}
\frac{dP}{dt} &= e_P((1-\phi_P)(f_{NP(H,N_l)} + f_{HP(H,N_l)}) + \phi_P I_P)P - \mu_P P \\
\frac{dN_a}{dt} &= m_N N_l - \mu_N N_a \\
\frac{dN_l}{dt} &= e_N((1-\phi_N)f_{HN(H)} + \phi_N I_N(N_l + N_a)) - (1-\phi_P)f_{NP(H,N_l)}P - (\mu_N + m_N)N_l \\
\frac{dH}{dt} &= \rho(K-H) - (1-\phi_N)f_{HN(H)}(N_l + N_a) - (1-\phi_P)f_{HP(H,N_l)}P
\end{aligned}
$$

##### 1.4.2 PBHF (IG Predator P = Predatory bug PB, IG prey N = Hoverfly HF)

$$
\begin{aligned}
\frac{dP}{dt} &= e_P((1-\phi_P)(f_{NP(H,N_l)} + f_{HP(H,N_l)}) + \phi_P I_P)P - \mu_P P \\
\frac{dN_a}{dt} &= m_N N_l - \mu_N N_a \\
\frac{dN_l}{dt} &= e_N(f_{(HN(H)}N_l + I_N N_a) - (1-\phi_P)f_{NP(H,N_l)}P - (\mu_N + m_N)N_l \\
\frac{dH}{dt} &= \rho(K-H) - f_{HN(H)}N_l - (1-\phi_P)f_{HP(H,N_l)}P
\end{aligned}
$$

##### 1.4.3 HFPB (IG Predator P = Hoverfly HF, IG prey N = Predatory bug PB)

$$\frac{dP_a}{dt} = m_P P_l - \mu_P P_a$$
$$\frac{dP_l}{dt} = e_P((f_{NP(H,N_l)} + f_{HP(H,N_l)})P_l + I_P P_a) - (\mu_P + m_P)P_l$$
$$\frac{dN_a}{dt} = m_N N_l - \mu_N N_a$$
$$\frac{dN_l}{dt} = e_N((1-\phi_N)f_{HN(H)} + \phi_N I_N (N_l + N_a)) - f_{NP(H,N_l)}P_l - (\mu_N + m_N)N_l$$
$$\frac{dH}{dt} = \rho(K - H) - (1-\phi_N)f_{HN(H)}(N_l + N_a) - f_{HP(H,N_l)}P_l$$

##### 1.4.4 HFHF (IG Predator P = Hoverfly HF, IG prey N = Hoverfly HF)

$$\frac{dP_a}{dt} = m_P P_l - \mu_P P_a$$
$$\frac{dP_l}{dt} = e_P((f_{NP(H,N_l)} + f_{HP(H,N_l)})P_l + I_P P_a) - (\mu_P + m_P)P_l$$
$$\frac{dN_a}{dt} = m_N N_l - \mu_N N_a$$
$$\frac{dN_l}{dt} = e_N(f_{(HN(H)}N_l + I_N N_a) - f_{NP(H,N_l)}P_l - (\mu_N + m_N)N_l$$
$$\frac{dH}{dt} = \rho(K - H) - f_{HN(H)}N_l - f_{HP(H,N_l)}P_l$$

#### 1.5 All the scenarios where the IG prey N was a Parasitoid

##### 1.5.1 LBPA (IG Predator P = Ladybird PB, IG prey N = Parasitoid PA)

When the IG prey N is a parasitoid we remove the natural death rate $\mu_N$ of the larva $N_l$ and only the adult $N_a$ fed on the herbivore $H$.

$$\frac{dP}{dt} = e_P(f_{NP(H,N_l)} + f_{HP(H,N_l)})P - \mu_P P$$
$$\frac{dN_a}{dt} = m_N N_l - \mu_N N_a$$
$$\frac{dN_l}{dt} = e_N f_{HN(H)}N_a - f_{NP(H,N_l)}P - m_N N_l$$
$$\frac{dH}{dt} = \rho(K - H) - f_{HN(H)}N_a - f_{HP(H,N_l)}P$$

##### 1.5.2 PBPA (IG Predator P = Predatory bug PB, IG prey N = Parasitoid PA)

$$\frac{dP}{dt} = e_P((1-\phi_P)(f_{NP(H,N_l)} + f_{HP(H,N_l)}) + \phi_P I_P)P - \mu_P P$$
$$\frac{dN_a}{dt} = m_N N_l - \mu_N N_a$$
$$\frac{dN_l}{dt} = e_N f_{HN(H)} N_a - (1-\phi_P) f_{NP(H,N_l)} P - m_N N_l$$
$$\frac{dH}{dt} = \rho(K-H) - f_{HN(H)} N_a - (1-\phi_P) f_{HP(H,N_l)} P$$

##### 1.5.3 HFPA (IG Predator P = Predatory bug PB, IG prey N = Parasitoid PA)

$$\frac{dP_a}{dt} = m_P P_l - \mu_P P_a$$
$$\frac{dP_l}{dt} = e_P((f_{NP(H,N_l)} + f_{HP(H,N_l)})P_l + I_P P_a) - (\mu_P + m_P)P_l$$
$$\frac{dN_a}{dt} = m_N N_l - \mu_N N_a$$
$$\frac{dN_l}{dt} = e_N f_{HN(H)} N_a - f_{NP(H,N_l)} P_l - m_N N_l$$
$$\frac{dH}{dt} = \rho(K-H) - f_{HN(H)} N_a - f_{HP(H,N_l)} P_l$$

#### 1.6 Functional responses

For all equations, the functional responses $f_{NP}$, $f_{HP}$ and $f_{HN}$ are the following (with changing values of $\phi_P$ and $\phi_N$ according to the predator type).

$$f_{NP(H,N_l)} = \frac{c_P(1-s)N_l}{h_P + (1-\phi_P)(sH + (1-s)N_l)}$$

$$f_{HP(H,N_l)} = \frac{c_P sH}{h_P + (1-\phi_P)(sH + (1-s)N_l)}$$

$$f_{HN(H)} = \frac{c_N H}{h_N + (1-\phi_N)H}$$

Where $C_P$ and $C_N$ are the maximum ingestion rates, $h_P$ and $h_N$ are the half saturation constants of IG predator P and IG prey N. When the IG predator P or IG prey N was a Predatory Bug we have $\phi_P = \phi_N = 0.5$. For all the other cases, $\phi_P = \phi_N = 0$

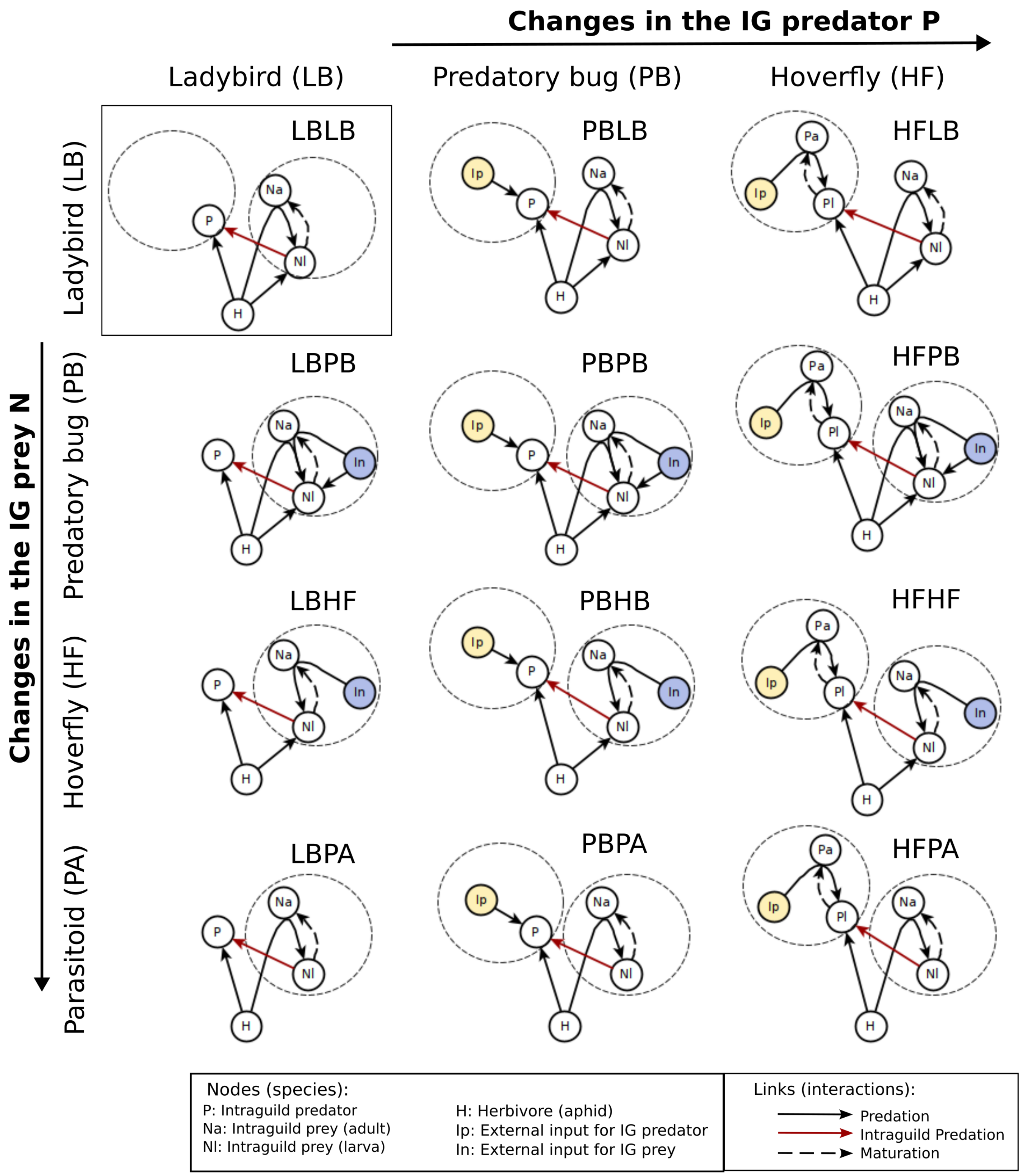


Figure S. 1: **12 IGP combinations** We represent the baseline **LBLB** module where both IG predator and IG prey belong to the LB type (square). Here the herbivore ($H$) can be eaten by the IG prey larva ($N_l$) and adult ($N_a$), as well as by the IG predator ($P$). Predation from $N_a$ directly transforms to biomass of the larva $N_l$. The IG predator also predates on the IG prey larva but not on the adult. Finally, $N_l$ matures into $N_a$. We then show the changes when other predator types acted either as an IG predator $P$ or as an IG prey $N$. The big dotted circles surround nodes that are different in relation to the baseline **LBLB** module. For the Predatory Bug type (PB) as an IG predator $P$ or as an IG prey $N$ we add external inputs $I_P$ (yellow) to $P$ or $I_N$ (blue) to both $N_a$ and $N_l$. In the case of the hoverfly type (HF), we divided the $P$ population into $P_a$ and $P_l$ to model how $I_P$ is consumed only by the adult $P_a$ when HF acted an IG predator, or only by $N_a$ when acted as a IG prey. For the parasitoid type (PA) as an IG prey, we assumed that the infected larva $N_l$ lies within the herbivore and is promoted by the infection of the $N_a$ of the herbivore $H$. The icons show some representative natural enemies of each predator type.

### 2 Additional methods: maximum external input and coexistence criteria

#### 2.1 Maximum external input

The maximum external input rates $I_N[max]$ and $I_P[max]$ were calculated as the value above which the predator involved (IG prey N or IG predator P) can live out only from this external source. This means that in the equilibrium, a small perturbation of this predator will grow, (or the derivative will be positive), given that the density of the herbivore and the other predator are zero. This value depended on the specific predator type.

##### 2.1.1 Predatory bug PB

For the predatory bug, given $\delta P > 0$, $\delta N_l > 0$, $\delta N_a > 0$ and $P^*$, $N_a^*$ and $N_l^*$ we use the jacobian (when needed) to solve:

$$\frac{d(P^* + \delta P)}{dt} > 0$$
$$\iff \quad I_P > \mu_P/\phi_P e_P = I_P[max] = 1.32$$

Equivalently, for the IG prey N:

$$\frac{d(N_a^* + \delta N_a)}{dt} > 0$$
$$\frac{d(N_l^* + \delta N_l)}{dt} > 0$$
$$\iff \quad I_N > \mu_N/\phi_N e_N = I_N[max] = 4.4$$

##### 2.1.2 Hoverfly PB

For the Hoverfly type, given $\delta P_a > 0$, $\delta P_l > 0$, $\delta N_l > 0$, $\delta N_a > 0$ and $P_a^*$,$P_l^*$, $N_a^*$ and $N_l^*$ the equilibria where only the external input is present, the criteria for persistence is calculated as:

$$\frac{d(P_a^* + \delta P_a)}{dt} > 0$$
$$\frac{d(P_l^* + \delta P_l)}{dt} > 0$$
$$\iff \quad I_P > \mu_P(m_P + \mu_P)/m_P e_P = I_P[max] = 1.7$$

And for the IG prey N:

Table S. 1: Values of external input rate used for the simulations in time units (t.u.)

| **Predator type** | **IG P or IG N** | **Maximum input rate** $I[max]$ **(t.u.)** | **Low input rate** ($10\%$) **(t.u.)** | **High input rate** ($90\%$)**(t.u.)** |
|---|---|---|---|---|
| Predatory Bug PB | IG predator P | $I_P[max] = 1.32$ | 0.13 | 1.2 |
| | IG prey N | $I_N[max] = 4.4$ | 0.4 | 4 |
| Hoverfly HF | IG predator P | $I_P[max] = 1.7$ | 0.18 | 1.6 |
| | IG prey N | $I_N[max] = 14.3$ | 1.4 | 13 |

$$\frac{d(N_a^* + \delta N_a)}{dt} > 0$$
$$\frac{d(N_l^* + \delta N_l)}{dt} > 0$$
$$\iff \quad I_N > \mu_N(m_N + \mu_N)/m_N e_N = I_N[max] = 14.3$$

For the simulations, we used $10$ and $90\%$ of the maximum external input rate values for $\phi = 0.5$ (Table S.1).

#### 2.2 Coexistence criteria

Polis and Holt (1989) have shown that in order to have coexistence between predators in an IGP module, the IG prey N must be a better competitor than the IG predator P for the herbivore H and reduce its density to relatively lower values. This is: $H_{[HP^*]} > H_{[HN^*]}$, which represent the value of the herbivore in the equilibria where only the IG predator or the IG prey are present. In the following, we write the expression of $H_{[HP^*]}$ and $H_{[HN^*]}$ for each predator type.

##### 2.2.1 Ladybird type LB

$$H_{[HN^*]} = \frac{h_N \mu_N}{e_N C_N - \mu_N} \tag{1}$$

$$H_{[HP^*]} = \frac{h_P \mu_P}{s(e_P C_P - \mu_P)} \tag{2}$$

##### 2.2.2 Predatory bug PB

$$H_{[HN^*]} = \frac{h_N \left( \frac{\mu_N + m_N}{e_N (1 + \frac{m_N}{\mu_N})} - \phi_N I_N \right)}{(1 - \phi_N) \left( C_N - \left( \frac{\mu_N + m_N}{e_N (1 + \frac{m_N}{\mu_N})} - \phi_N I_N \right) \right)} \tag{3}$$

$$H_{[HP^*]} = \frac{h_P \left( \frac{\mu_P}{e_P} - \phi_P I_P \right)}{(1 - \phi_P) s \left( C_P - \left( \frac{\mu_P}{e_P} - \phi_P I_P \right) \right)} \tag{4}$$

###### 2.2.3 Hoverfly HF

$$H_{[HN^*]} = \frac{h_N\left(\frac{\mu_N+m_N}{e_N} - \frac{I_N m_N}{\mu_N}\right)}{C_N - \left(\frac{\mu_N+m_N}{e_N} - \frac{I_N m_N}{\mu_N}\right)} \tag{5}$$

$$H_{[HP^*]} = \frac{h_P\left(\frac{m_P+\mu_P}{e_P} - \frac{I_P m_P}{\mu_P}\right)}{s\left(C_P - \left(\frac{m_P+\mu_P}{e_P} - \frac{I_P m_P}{\mu_P}\right)\right)} \tag{6}$$

###### 2.2.4 Parasitoid PA

$$H_{[HN^*]} = \frac{h_N \mu_N}{e_N C_N - \mu_N} \tag{7}$$

So for each of these models we have to check the following inequality as the coexistence criteria.

$$H_{[HN^*]} < H_{[HP^*]} \tag{8}$$

For all predator types, this inequality depend on the IGP symmetry $s$ and on the external inputs $I_N$ and $I_P$, which we varied within the models, but do not depend on the productivity $K$. So, in order to see how these parameters affected coexistence, we plot the "maximum $s$ value" above which the coexistence criteria is not met (giving the fixed parameters used in the main text) (Fig. 2). This means that below this value coexistence is just *possible* and that above it, coexistence is not possible. We observed that for models without external input (**LBLB** and **LBPA**) the maximum value for $s$ equals to 1. This means that for $s < 1$ coexistence is possible. Adding an external value to the IG prey N ($I_N$) does not really affect this value, except for low values for the **LBPB** scenario (where $s < 0.75$). Adding a high external value to the IG predator P ($I_P$) drops the value of $s$ for the **PBLB**, **HFLB**, **PBPA** and **HFPA** modules ($s < 0.2$). When both external inputs were present (**PBPB**,**HFHF**,**PBHF**,**HFPB**), a low ($I_N$) and a high ($I_N$) drop the value of $s$ ($s < 0.2$). However, when the external input for the IG prey ($I_N$) is high, the $s$ value can go up to 1.

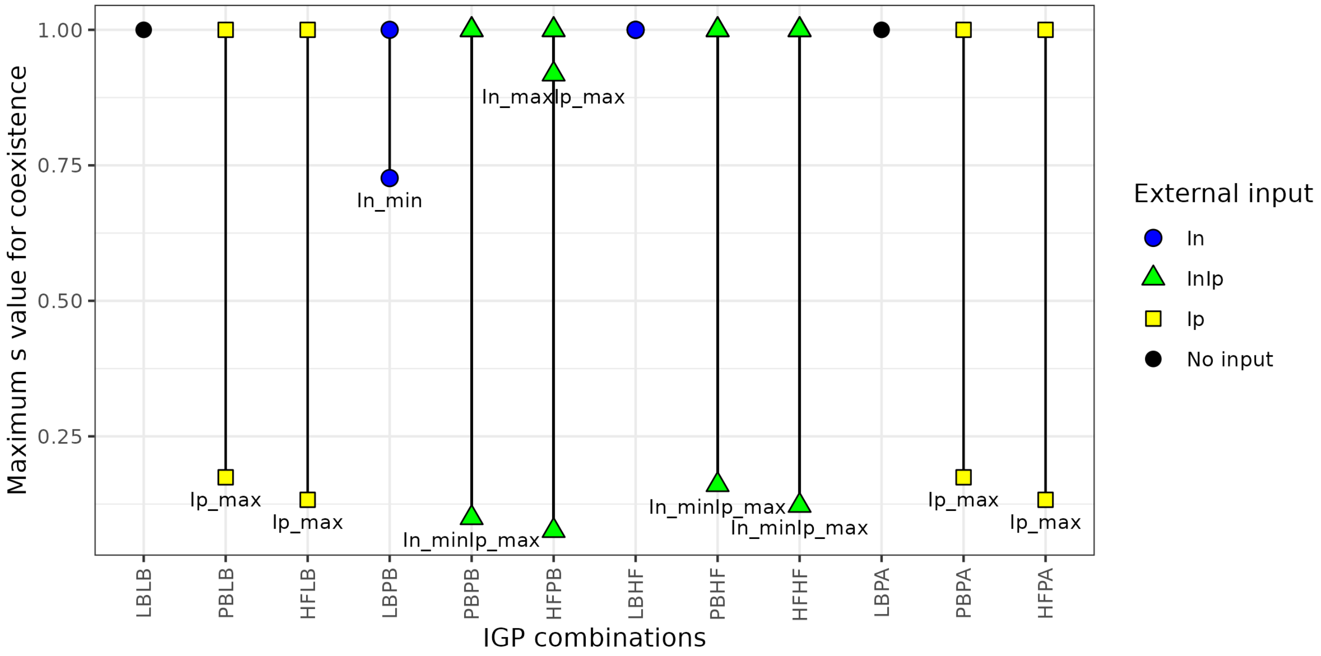


Figure S. 2: **Role of the IGP symmetry s for the criteria of coexistence for the different IGP combinations**. For each IGP module we show the maximum $s$ value" above which coexistence is not theoretically possible. The shapes and colors represent the external inputs for each scenario. We labeled those in which $s < 1$. The line represents how the value of the external input affected the range of IGP symmetry $s$.

### 3 Additional results: boundary curves of the LBLB module

The herbivore-only equilibrium is stable as long as the population growth rates of IG prey and IG predator are negative at a particular equilibrium herbivore density. Since IG preys have lower herbivore requirements than IG predators ($H_{[HP^*]} > H_{[HN^*]}$), a small perturbation of $N$ can invade the herbivore-only equilibrium given that the following threshold $b_N$ is positive:

$$b_N(K) = \frac{e_N C_N K}{h_N + K} - \mu_N \tag{9}$$

which depends only on the productivity of the system (in the absence of $P$, the consumption of $H$ by $N$ is not affected by $s$). $b_N(K) > 0$ means that the biomass derived by the consumption of the herbivore (which density is at equilibrium) must overcome its death rate. Similarly, IG predators $P$ can invade a herbivore-only equilibrium if $b_P > 0$:

$$b_P(K,s) = \frac{e_P C_P s K}{h_P + sK} - \mu_P \tag{10}$$

This criteria will not be relevant since for every $S \in [0,1]$ and $K > 0$, if $b_P > 0$ then $b_N > 0$. Two other boundary curves are derived if we ask the necessary conditions for the $N$ and $P$ to invade the equilibrium when the other predator is already present. $P$ can invade $HN^*$ equilibrium if $b_{N\leftarrow P} > 0$:

$$b_{N\leftarrow P}(K,s) = \frac{e_P C_P (sH_{[HN^*]} + (1-s)Nl_{[HN^*]})}{h_P + sH_{[HN^*]} + (1-s)Nl_{[HN^*]}} - \mu_P \tag{11}$$

where $H_{[HN^*]} = \mu_N h_N/(e_N C_N - \mu_N)$ and $Nl_{[HN^*]} = e_N \rho(K - H_{[HN^*]})/(\mu_N + m_N)$ are the values of the herbivore and the larval stage of $N$ in the $HN^*$ equilibrium. This curve bends with higher values of $s$. So for low $s$ (near tritrophic), medium productivities are sufficient for $P$ to invade the $HN^*$ equilibrium (as it mostly feeds on $N$. But as soon as the network becomes more competitive ($s \to 1$), $P$ need higher amounts of herbivore to overcome the competition with $N$ and invade the system. Finally, N can still invade $HP^*$ equilibrium if $b_{P\leftarrow N} > 0$.

$$b_{P\leftarrow N}(K,s) = \frac{\mu_N + m_N}{\mu_N}\left(\frac{e_N C_N H_{[HP^*]}}{h_N + H_{[HP^*]}} - \mu_N\right) - \frac{C_P(1-s)e_P\rho(K - H_{[HP^*]})}{\mu_P(h_P + sH_{[HP^*]})} \tag{12}$$

where $H_{[HP^*]} = \mu_P h_P / s(e_P C_P - \mu_P)$ is the value of the herbivore in the $HP^*$ equilibrium. Interestingly, this curve is bent both a low and high values of $s$. At low $s$, $P$ relies a lot on $N$, so the system would need to reach very high productivities for $N$ to be able to invade a $HP^*$ equilibrium. At high $s$, $P$ mostly feeds on the herbivore, but as $N$ is a better competitor for it, we would need a lot of herbivore for $N$ not being able to invade. The crossings of these curves define the regions for ecological invasibility and determine the different equilibria. Given some $(s,K)$ that satisfies $b_N > 0$, we have different scenarios. If $(s,K)$ fulfills only one of the two predator-herbivore invasibility criteria ($b_{N\leftarrow P} > 0$ or $b_{P\leftarrow N} > 0$) we obtain a $HN^*$ or a $HP^*$ equilibria. If $(s,K)$ does not satisfy any of these criteria, we have a zone of bistability of $HN^*$ and $HP^*$ equilibria whose final state depends on the initial conditions. Now, if $(s,K)$ satisfies a "mutual invasibility criteria" ($b_{N\leftarrow P} > 0$ and $b_{P\leftarrow N} > 0$) then we have an equilibrium zone of $HNP^*$. Given a fixed value of $s$, this range of $K$ for which both predators can invade the other only exists if the minimum value of $K$ need for the IG predator $P$ to invade a $HN$ system (the value of $K$ at which $b_{N\leftarrow P} > 0$, this is $K_{N\leftarrow P}$) is lower than the maximum value of $K$ above which $N$ can not longer invade a $HP$ system (the value of $K$ at which $b_{P\leftarrow N} > 0$, or $K_{P\leftarrow N}$)). However, we also have a zone of bistability of $HP^*$ and $HNP^*$ where only $b_{N\leftarrow P} > 0$ is satisfied (and should correspond only to $HP^*$ equilibrium). In addition, with very low levels of $s$ and high levels of $K$, the system exhibits a Hopf Bifurcation above the which the $HNP^*$ becomes periodic.

### 4 Additional results: on the preference $\phi$

We evaluated the role that the parameter $\phi$ played on the area of the region of coexistence for the cases where the Predatory Bug PB was present, either as an IG predator P or as an IG prey N. For this, we ran the **LBPB** and **PBLB** modules using three values of $\phi_N$ and $\phi_P$ $(0.2, 0.5, 0.8)$. For each of these values, the maximum external input also changed to satisfy that the system is an IGP system, and the low and high values of $I$ accordingly (Table S.2.). We observe that the general effect of increasing the external input is maintained independently of the values of the preference. The scenario **PBPB** was not modeled as it is a combination of the last two.

Table S. 2: Values of external input rate when changing the preference for the Predatory Bug PB (in time units (t.u.))

| **Predator type** | Preference ($\phi$) | **IG P or IG N** | **Maximum input rate** $I[max]$ **(t.u.)** | **Low input rate** ($10\%$) **(t.u.)** | **High input rate** ($90\%$)**(t.u.)** |
|---|---|---|---|---|---|
| Predatory Bug PB | 0.2 | IG predator P | $I_P[max] = 3.3$ | 0.33 | 3.0 |
| | | IG prey N | $I_N[max] = 11$ | 1.1 | 9.9 |
| | 0.8 | IG predator P | $I_P[max] = 0.83$ | 0.083 | 0.75 |
| | | IG prey N | $I_N[max] = 2.8$ | 0.28 | 2.5 |

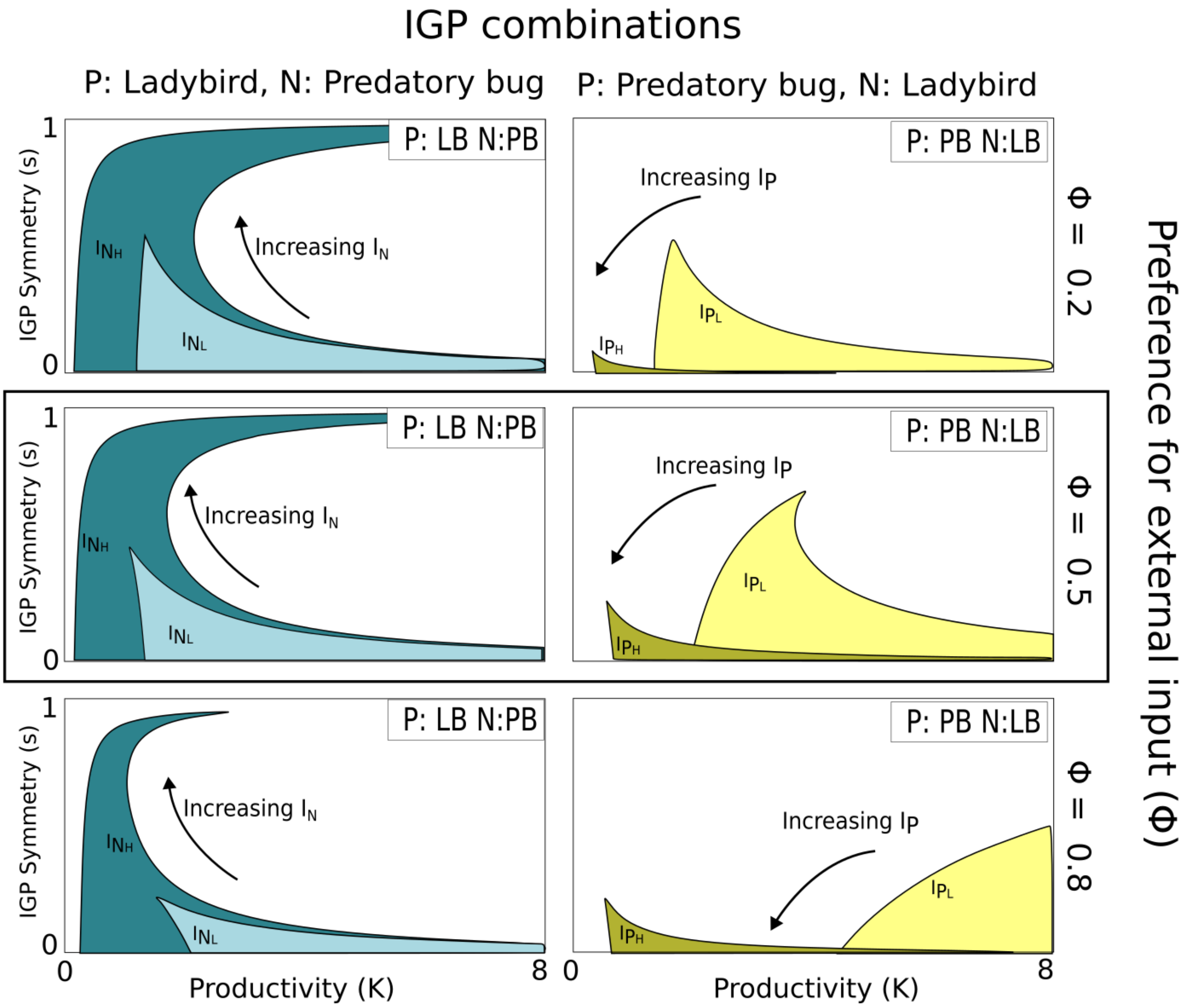


Figure S. 3: **Effect on the preferences $\phi_N$ and $\phi_P$ on the change of the area of coexistence with different external inputs.** The values of $\phi_N$ (**LBPB** modules) and $\phi_P$ (**PBLB** modules) where $0.2, 0.5, 0.8$. For each of these we used different values of $I_N$ and $I_P$ respectively (see Table S.2.

### 5 Additional results on herbivore suppression

We plot in this section, the IG prey N and the IG predator P densities (Figures S.4 and S.5) for three different values of IGP symmetries across the productivity. We also show the expanded Table S.3 for the mean coexistence and herbivore density of each IGP combination and its relative increase or decrease in comparison to the baseline **LBLB**.

Table S. 3: **Influence of different predator types on the coexistence and herbivore suppression**. For each IGP module category, range $K$ (Low: $K < 5$, High: $K \geq 5$ and IGP symmetry $s$, we calculated the mean herbivore density and the mean coexistence.

| IG Predator P | IG prey N | Ip | In | IGP symmetry (s) | Productivity (K) | Mean H | Mean Coexistence |
|---|---|---|---|---|---|---|---|
| LB | LB | 0 | 0 | Tritrophic ($s = 0.1$) | Low | 0.98 | 0.75 |
| | | | | | High | 3.59 | 0.00 |
| | | | | Symmetric ($s = 0.5$) | Low | 0.61 | 0.00 |
| | | | | | High | 0.72 | 0.00 |
| | | | | Competitive ($s = 0.9$) | Low | 0.34 | 0.00 |
| | | | | | High | 0.40 | 0.00 |
| PB/HF | LB | Low $I_P$ | 0 | Tritrophic ($s = 0.1$) | Low | 0.42 | 0.40 |
| | | | | | High | 2.48 | 1.00 |
| | | | | Symmetric ($s = 0.5$) | Low | 0.54 | 0.40 |
| | | | | | High | 1.24 | 0.00 |
| | | | | Competitive ($s = 0.9$) | Low | 0.28 | 0.00 |
| | | | | | High | 0.28 | 0.00 |
| LB | PB/HF | 0 | High $I_N$ | Tritrophic ($s = 0.1$) | Low | 1.14 | 1.00 |
| | | | | | High | 3.56 | 0.25 |
| | | | | Symmetric ($s = 0.5$) | Low | 0.56 | 0.38 |
| | | | | | High | 0.72 | 0.00 |
| | | | | Competitive ($s = 0.9$) | Low | 0.14 | 0.88 |
| | | | | | High | 0.38 | 0.13 |
| PB/HF | PB/HF | Low $I_P$ | High $I_N$ | Tritrophic ($s = 0.1$) | Low | 0.59 | 1.00 |
| | | | | | High | 2.87 | 1.00 |
| | | | | Symmetric ($s = 0.5$) | Low | 0.43 | 0.88 |
| | | | | | High | 1.24 | 0.00 |
| | | | | Competitive ($s = 0.9$) | Low | 0.06 | 0.75 |
| | | | | | High | 0.19 | 1.00 |
| LB | PA | 0 | 0 | Tritrophic ($s = 0.1$) | Low | 2.20 | 0.83 |
| | | | | | High | 3.59 | 0.00 |
| | | | | Symmetric ($s = 0.5$) | Low | 0.72 | 0.00 |
| | | | | | High | 0.72 | 0.00 |
| | | | | Competitive ($s = 0.9$) | Low | 0.40 | 0.00 |
| | | | | | High | 0.40 | 0.00 |
| PB/HF | PA | Low $I_P$ | 0 | Tritrophic ($s = 0.1$) | Low | 2.23 | 1.00 |
| | | | | | High | 5.40 | 0.67 |
| | | | | Symmetric ($s = 0.5$) | Low | 1.06 | 0.25 |
| | | | | | High | 1.24 | 0.00 |
| | | | | Competitive ($s = 0.9$) | Low | 0.56 | 0.20 |
| | | | | | High | 0.69 | 0.00 |

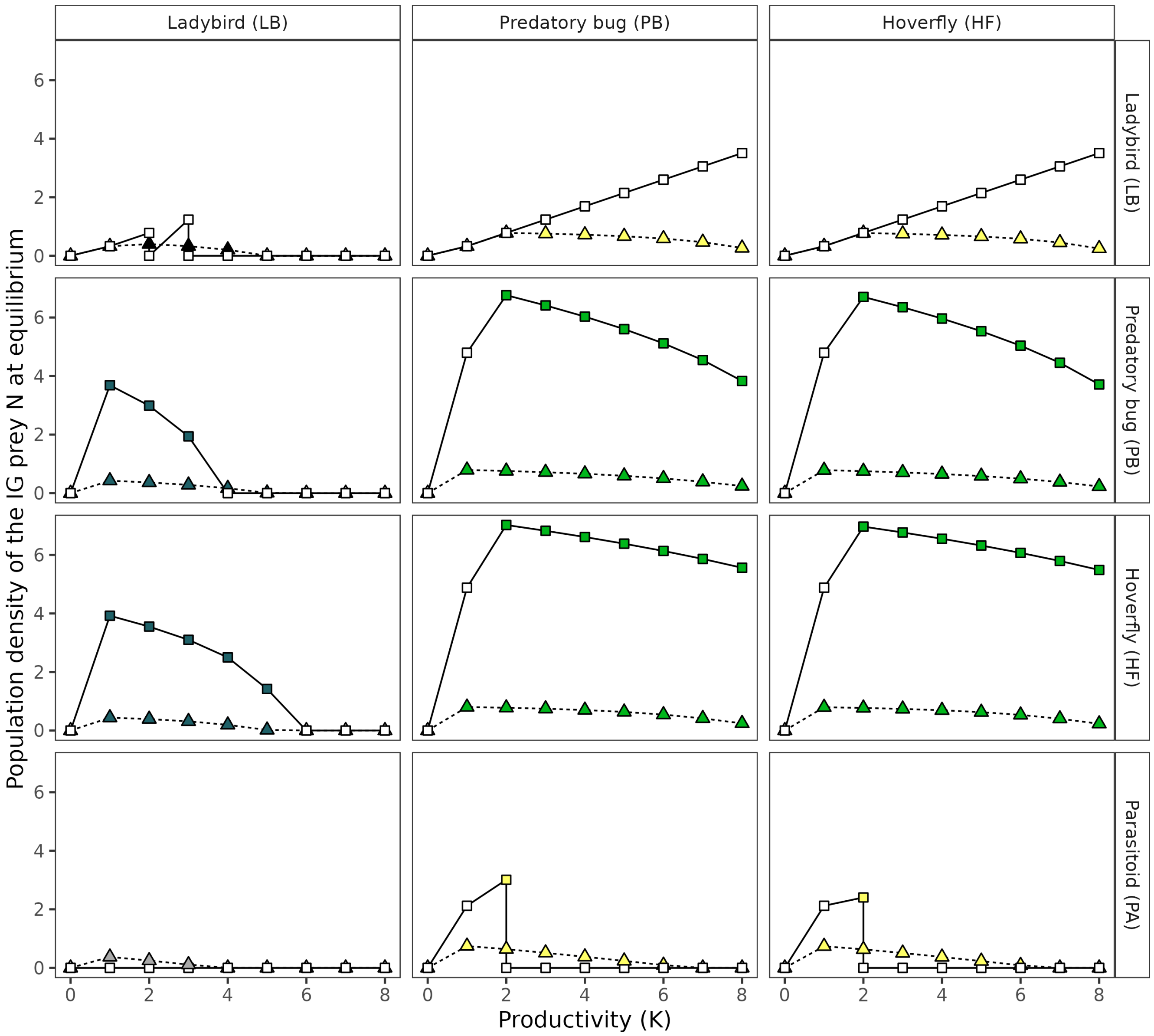


Figure S. 4: **IG prey in the regions of coexistence**. We show the population density of the IG prey N at equilibrium for each of the 12 IGP modules, along the productivity gradient and for three representative IGP symmetries: tritrophic-like ($s = 0.1$, triangles and dashed lines), symmetric ($s = 0.5$, circles and short-dashed line) and competitive-like ($s = 0.9$, squares and continuous line). We represent if the system was at a coexistence equilibria ($HNP^*$ equilibrium, filled shapes) or not (without distinguish between $H^*$, $HN^*$ or $HP^*$ equilibria, empty shapes). For the IGP modules that include the predatory bug (PB) or the hoverfly (HF) types as IG predators or IG preys, we include only the values of external input $I_P = I_{P_L}$ and $I_N = I_{N_H}$ that increased coexistence. In this sense, the colors of the filled circles follow the code color of figure 4 ($I_{N_H}$: dark blue, $I_{N_L}$: light blue, $I_{P_L}$: yellow, $I_{P_L}I_{N_L}$: vivid green, $I_{P_L}I_{N_H}$: green, **LBPA**: gray, **LBLB**: black).

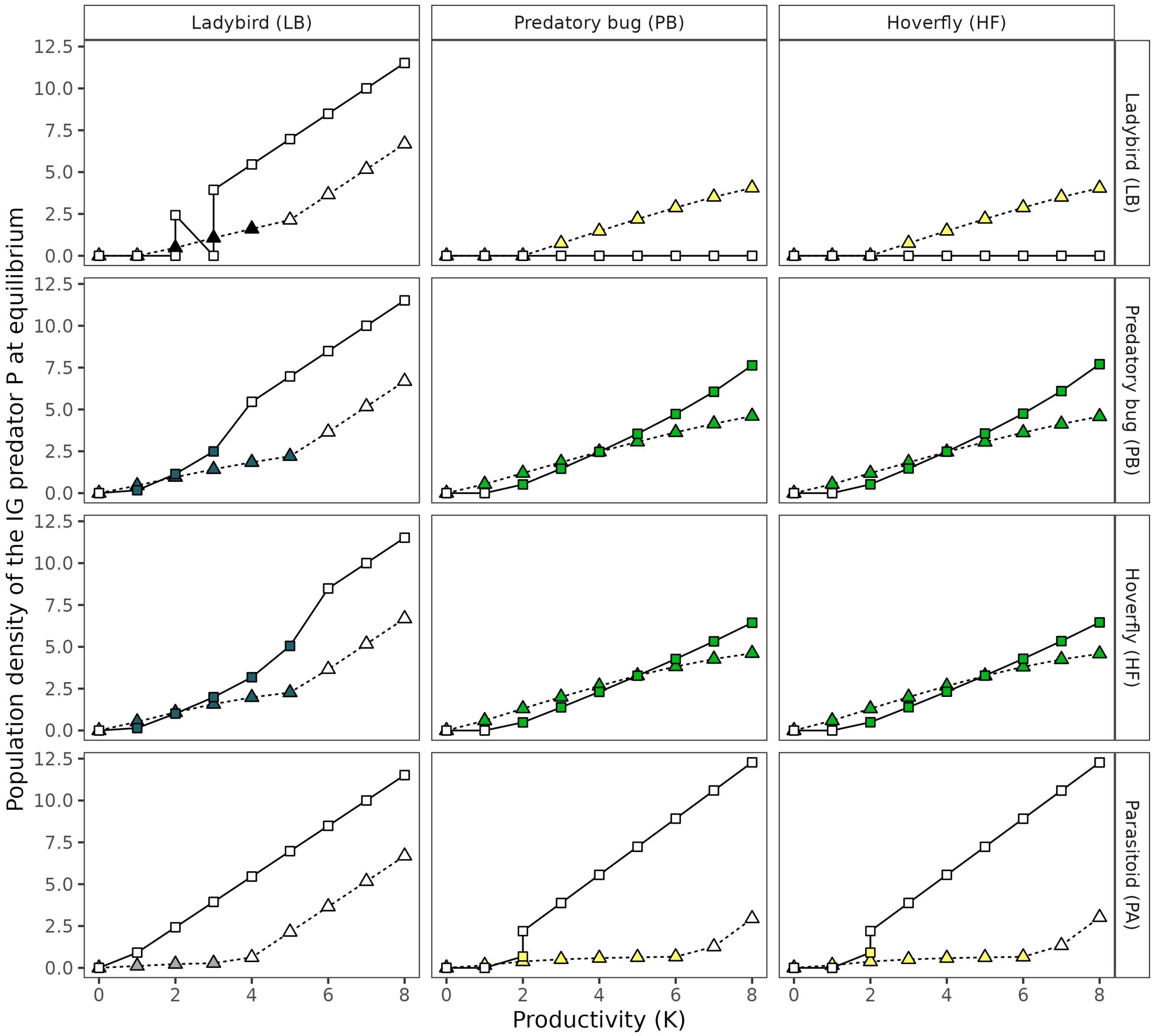


Figure S. 5: **IG predator in the regions of coexistence**. We show the population density of the IG predator P at equilibrium for each of the 12 IGP modules, along the productivity gradient and for three representative IGP symmetries: tritrophic-like ($s = 0.1$, triangles and dashed lines), symmetric ($s = 0.5$, circles and short-dashed line) and competitive-like ($s = 0.9$, squares and continuous line). We represent if the system was at a coexistence equilibria ($HNP^*$ equilibrium, filled shapes) or not (without distinguish between $H^*$, $HN^*$ or $HP^*$ equilibria, empty shapes). For the IGP modules that include the predatory bug (PB) or the hoverfly (HF) types as IG predators or IG preys, we include only the values of external input $I_P = I_{P_L}$ and $I_N = I_{N_H}$ that increased coexistence. In this sense, the colors of the filled circles follow the code color of figure 4 ($I_{N_H}$: dark blue, $I_{N_L}$: light blue, $I_{P_L}$: yellow, $I_{P_L}I_{N_L}$: vivid green, $I_{P_L}I_{N_H}$: green, **LBPA**: gray, **LBLB**: black).
